# Evolutionary and biomedical implications of sex differences in the primate brain transcriptome

**DOI:** 10.1101/2022.10.03.510711

**Authors:** Alex R. DeCasien, Kenneth L. Chiou, Camille Testard, Arianne Mercer, Josué E. Negrón-Del Valle, Samuel E. Bauman Surratt, Olga González, Michala K. Stock, Angelina V. Ruiz-Lambides, Melween I. Martinez, Cayo Biobank Research Unit, Susan C. Antón, Christopher S. Walker, Jérôme Sallet, Melissa A. Wilson, Lauren J. N. Brent, Michael J. Montague, Chet C. Sherwood, Michael L. Platt, James P. Higham, Noah Snyder-Mackler

**Author notes:** These authors contributed equally to this manuscript (Alex R. DeCasien, Kenneth L. Chiou). These authors are co-senior authors (James P. Higham, Noah Snyder-Mackler).

## Abstract

Humans exhibit sex differences in the prevalence of many neurodevelopmental and neurodegenerative conditions. To better understand the translatability of a critical nonhuman primate model, the rhesus macaque, we generated one of the largest multibrain region bulk transcriptional datasets for this species and characterized sex-biased gene expression patterns. We demonstrate that these patterns are similar to those in humans and are associated with overlapping regulatory mechanisms, biological processes, and genes implicated in sex-biased human disorders, including autism. We also show that sex-biased genes exhibit greater genetic variance for expression and more tissue-specific expression patterns, which may facilitate the rapid evolution of sex-biased genes. Our findings provide insights into the biological mechanisms underlying sex-biased disease and validate the rhesus macaque model for the study of these conditions.

## Introduction

Humans exhibit sex* differences in prevalence, presentation, and progression of many psychiatric, neurodevelopmental, and neurodegenerative conditions (*see Note 1). For example, depression (1), anxiety (2) and Alzheimer’s disease (AD) (3) are more prevalent in females, whereas attention deficit hyperactivity disorder (ADHD) (4), autism spectrum disorders (ASD) (5), schizophrenia (6), and Parkinson’s disease (7) occur more often in males. Although gender-biases in the applicability of diagnostic criteria certainly contribute to these differences (e.g., 8), neurobiological sex differences are likely to play a critical role, as multiple diagnostically-distinct disorders show the same sex bias during the same developmental window (e.g., male-biased early-onset neurodevelopmental disorders) and sex-biased disorders tend to emerge during particularly dynamic neurodevelopmental periods that involve changes to sex hormone concentrations (e.g., adolescence, menopause) (9, 10). Studies of post-mortem human brains have highlighted a molecular mechanism that may underlie such differences: many genes associated with these conditions are also expressed at different levels in healthy male and female brains (11–19). However, our understanding of the proximate and evolutionary sources of normative transcriptomic sex differences in the human brain is currently limited due to: i) a dearth of post-mortem human tissues, which tend to be very heterogeneous in terms of co-occurring diseases and processing methods; and ii) the fact that most work on neurobiological sex differences has been conducted on laboratory rodents, which are distantly related to and neuroanatomically distinct from humans.

Among existing animal models, rhesus macaques arguably have the greatest translatability to humans due to their close evolutionary relatedness, overall similar biology, wide availability, and deep knowledge acquired through over a century of cumulative biological and behavioral study. Like humans, macaques have primate-specific prefrontal cortical areas implicated in multiple neurological disorders (20), exhibit complex social behaviors that are mediated by similar neural circuits (21), and undergo extended brain development (relative to smaller model species such as rodents and marmosets, which are highly altricial) (22). Given that species-specific evolutionary mechanisms (e.g., mate choice, mate competition, parental care) (23) can produce species-specific sex differences in behavior and neurobiology, it is critical to explicitly test whether model organisms also exhibit human-like brain sex differences. This may be particularly relevant to the brain transcriptome, as sex-biased gene expression patterns tend to be species-specific (24), and sex-biased genes evolve faster than non-sex-biased genes in terms of changes to both coding sequence and gene expression (25–31). In cases where model species do exhibit human-like sex-biased associations with molecular signatures of disease, these conserved mechanisms may inform our understanding of sex-linked biological versus cultural drivers of sex-biased human diseases. Previous transcriptomic studies of the rhesus macaque brain (32–36) did not focus on sex differences and/or had restricted sampling of individuals, limiting our understanding of the extent to which humans and rhesus macaques share sex differences in brain gene expression, or of the evolutionary mechanisms that may have contributed to any differences. Filling in these gaps regarding the critically important rhesus macaque model could greatly aid in the development of potential therapies for sex-biased brain disorders in humans.

To address this, we generated one of the largest nonhuman primate brain transcriptional datasets (n=527 samples), and quantified sex differences in gene expression across 15 brain regions (Figure 1; Supplementary Table 1) from 36 free-ranging adults (20 females, 16 males; identified using chromosomal and phenotypic sex; Supplementary Figure 1; Supplementary Table 2). This substantial sample size allowed us to characterize, for the first time, patterns of sex-biased gene expression across the rhesus macaque brain, to link these patterns to human sex differences in the brain and disease, and to illuminate some of the evolutionary mechanisms underlying these patterns.

**Figure 1.**
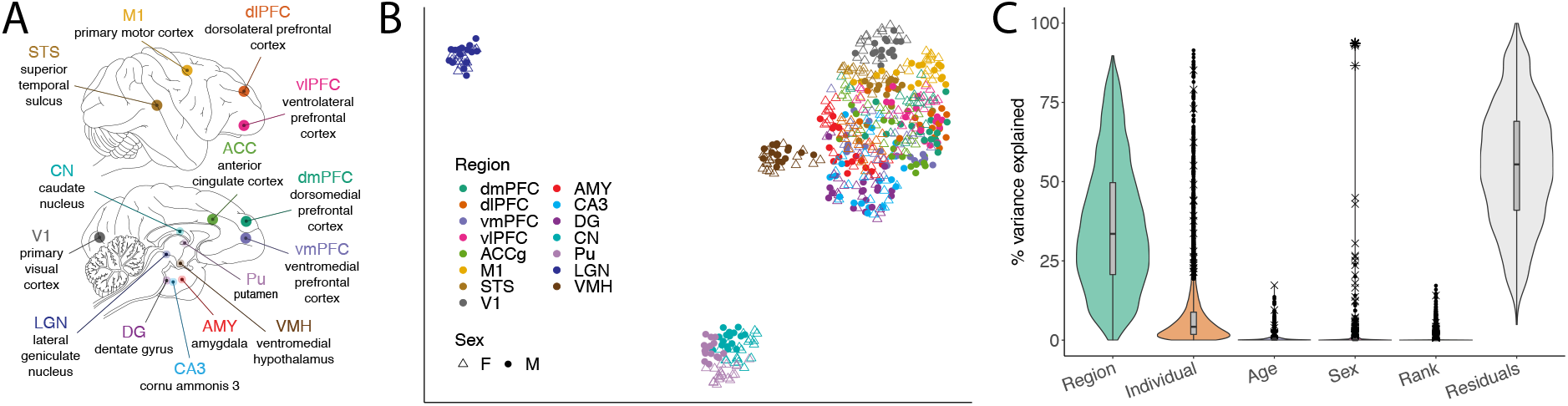
Experimental design and global expression patterns. (**A**) 15 brain regions sampled in the current study. Top = lateral view. Bottom = medial view. Some structures are internal and cannot be viewed from the planes depicted. (**B**) Uniform Manifold Approximation and Projection (UMAP) plot of expression data. Each point represents one sample (N=527). Colors indicate region and shape indicates sex (see legend). (**C**) Violin plots with overlaid boxplots of variance proportions for each gene and variable from variance partitioning analysis. Boxplots indicate the median (black horizontal line), first and third quartiles (i.e., interquartile range, IQR; lower and upper hinges), and ranges extending from each to 1.5 x IQR beyond each hinge (whiskers). Points represent individual genes that are outliers (i.e., beyond whiskers), and their shape indicates the chromosomal location (autosome = •, X chromosome = ×, Y chromosome = *).

## Results

### Sex-biased gene expression is largely shared across brain regions

We first quantified the drivers of global gene expression variation across all 527 samples for 12,672 detectably expressed genes in 15 regions (8 cortical regions, 2 hippocampal subregions, 2 striatal subregions, amygdala, hypothalamus, thalamus; Methods; Figure 1; Supplementary Table 1). As expected, the primary driver of variance in brain gene expression was the sampled brain region (mean = 36.12%; Figure 1; Supplementary Table 3), likely due to regional differences in cell composition and function. Indeed, regions in topographical proximity and with functional overlap exhibited more similar transcriptional profiles (Figure 1, Supplementary Figures 2-3). Although demographic and behavioral factors explained much less variation in the expression of individual genes across the whole brain (means: sex = 0.50%, dominance rank = 0.41%, age = 0.49%; Figure 1; Supplementary Table 3), their explanatory power was slightly higher within regions, particularly for age (means: sex = 0.78%, dominance rank = 1.46%, age = 3.66%; Supplementary Table 4). Sex explained significantly more variance for genes located on sex chromosomes compared to autosomal genes (overall brain means: Y chromosome = 92.74%, X chromosome = 1.16%, autosomes = 0.41%; Tukey’s HSD p_adj_ < 0.001; Figure 1; Supplementary Table 3).

**Figure 2.**
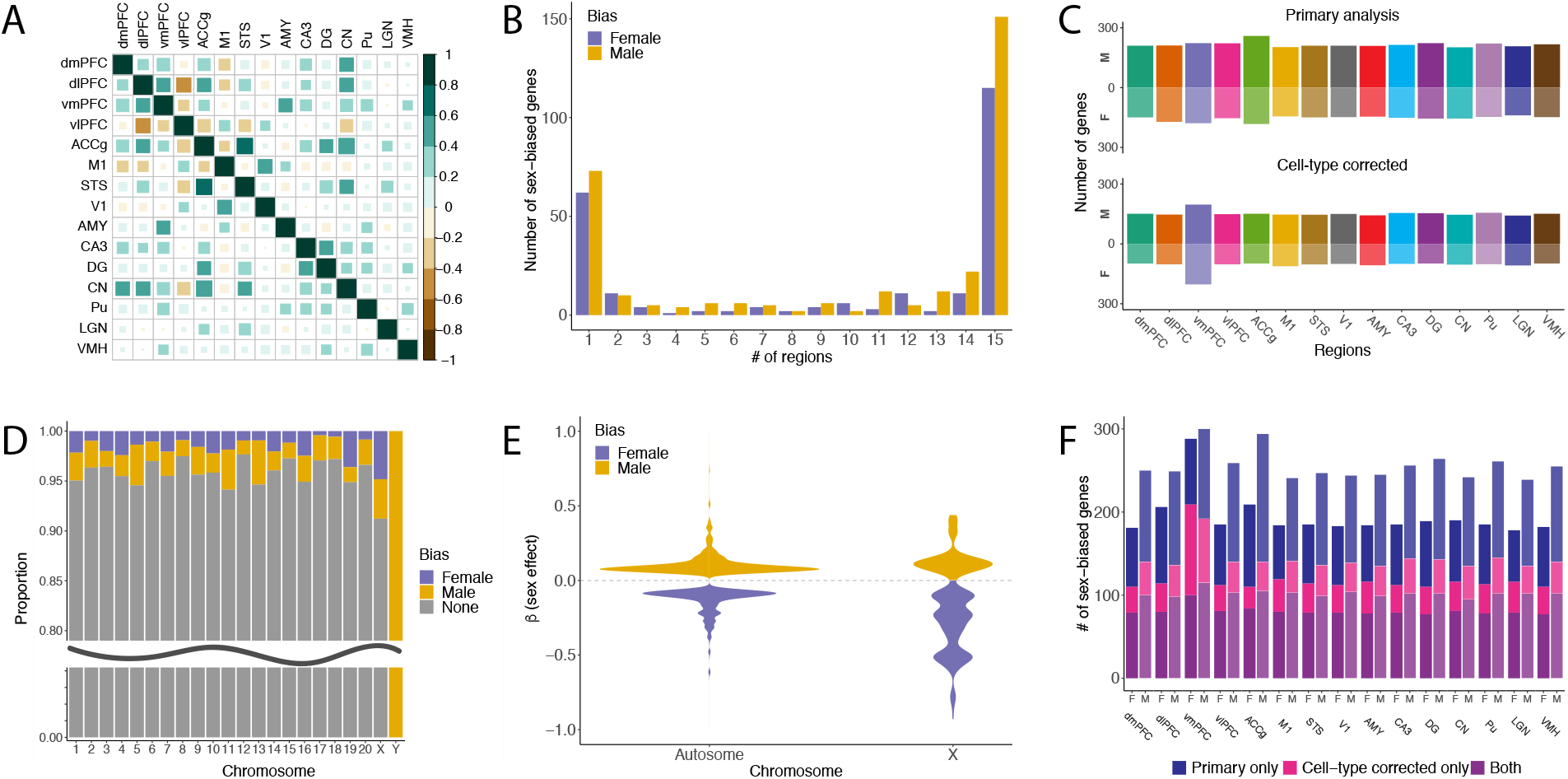
Regional, chromosomal, and cell-type distributions of sex-biased genes. (**A**) Correlation plot for pre-mashr sex effect sizes (from EMMREML) across regions (Spearman’s ρ). Teal = positive correlation, brown = negative correlation, size of square indicates strength of correlation. Of these inter-regional correlations, 77 are significantly positive, 23 are significantly negative, and are 5 not significant (p > 0.05). (**B**) Bar chart of the number of sex-biased genes (LFSR < 0.05) shared across different numbers of regions identified by our primary mashr analyses. (**C**) Counts of sex-biased genes identified by mashr (LFSR < 0.05) using un-adjusted (top) and cell type-corrected (bottom) expression data. M = male-biased, F = female-biased. (**D**) Proportions of genes on each chromosome that are not biased in any region (grey), female-biased in at least 1 region (purple), or male-biased in at least 1 region (yellow). The sex chromosomes are significantly enriched for sex-biased genes. (**E**) Violin plots of sex effect sizes (mashr betas) for sex-biased autosomal versus X chromosome genes. (**F**) Stacked bar plots of the number of male- and female-biased genes identified per region in our primary and/or cell-type corrected analyses.

Next, we estimated sex biases in gene expression within each brain region using linear mixed models controlling for age, dominance rank, technical covariates, and genetic relatedness (37). Sex effects were similar across regions, such that genes more highly expressed in females in one region tended to also be more highly expressed in females in all other regions (Figure 2; Supplementary Figure 4). This is consistent with observations of shared sex effects across other tissues in multiple species and suggests shared gene regulation across functionally and cellularly distinct tissues (24). We then implemented multivariate shrinkage (38) across regions to increase power, improve precision of our sex effect estimates, and estimate local false sign rates (LFSRs) (39). LFSRs quantify our confidence in the direction of effect estimates and are more conservative than the local false discovery rates (LFDRs), which instead measure the confidence that the effect is non-zero (39). In total, 4.4% (561/12,672) of genes expressed in the brain were differentially expressed between males and females (LFSR < 0.05) in at least one region (Figure 2; Supplementary Table 5), similar to human studies (average across 8 overlapping brain tissues = 6.5%) (12). Most sex-biased genes exhibited significant sex differences in the same direction in a majority of regions (66.8% were biased in at least 8/15 tissues; Figure 2), consistent with shared regulatory mechanisms. Of the identified sex-biased genes, 7.1% were located on the X chromosome, 1.6% on the Y chromosome, and 91.3% on autosomes (Figure 2). For all regions, the number of male-biased genes (i.e., genes more highly expressed in males relative to females) was higher than the number of female-biased genes (across all sex-bias genes: male-biased = 57%, female-biased = 43%; Figure 2); however, female-biased genes exhibited significantly stronger sex effects (mean |β| = 0.16) than male-biased genes (mean |β| = 0.12; t-test: p < 5.2e-36; excluding N = 9 Y chromosome genes, which are not expressed in females). In particular, female-biased X chromosome genes exhibited significantly larger sex effects (N = 22 genes; mean |β| = 0.32) than either male-biased X chromosome genes (N = 18; mean |β| = 0.15; Tukey’s HSD p_adj_ < 0.001) or sex-biased autosomal genes (female: N = 218, mean |β| = 0.14, p_adj_ < 0.001; male: N = 294: mean |β| = 0.11, p_adj_ < 0.001) (Figure 2), which is likely to reflect genes that escape X chromosome inactivation (XCI). Notably, all female-biased X-linked genes that we identified in macaques are known XCI escapees in humans (12), which may suggest conserved patterns of XCI escape across species. Other model organisms, including mice, do not exhibit these patterns due to a relatively low rate of XCI escape (40), which may further limit their translatability for sex-linked human conditions.

### Sex-biased brain gene expression is similar in rhesus macaques and humans

To investigate whether humans and rhesus macaques exhibit similar sex differences in brain gene expression, we compared estimated sex effects from this study (described above) to those from an analysis of the human GTEx data (V8) for 8 overlapping brain regions (controlling for age and technical effects; see Methods). Similar to our findings in macaques, sex explained a mean 0.49% of the variation in gene expression across the human GTEx brain samples. We also found that sex effects were similar across species (for all non-Y chromosome genes across all 8 regions: median Spearman’s correlation (ρ) = 0.16 [0.14-0.21]; p_adj_ < 0.05) Figure 3; Supplementary Figure 5; Supplementary Table 6). Together, these observations suggest that global transcriptomic sex differences in macaque and human brains largely mirror one another across both cortical and subcortical brain regions.

**Figure 3.**
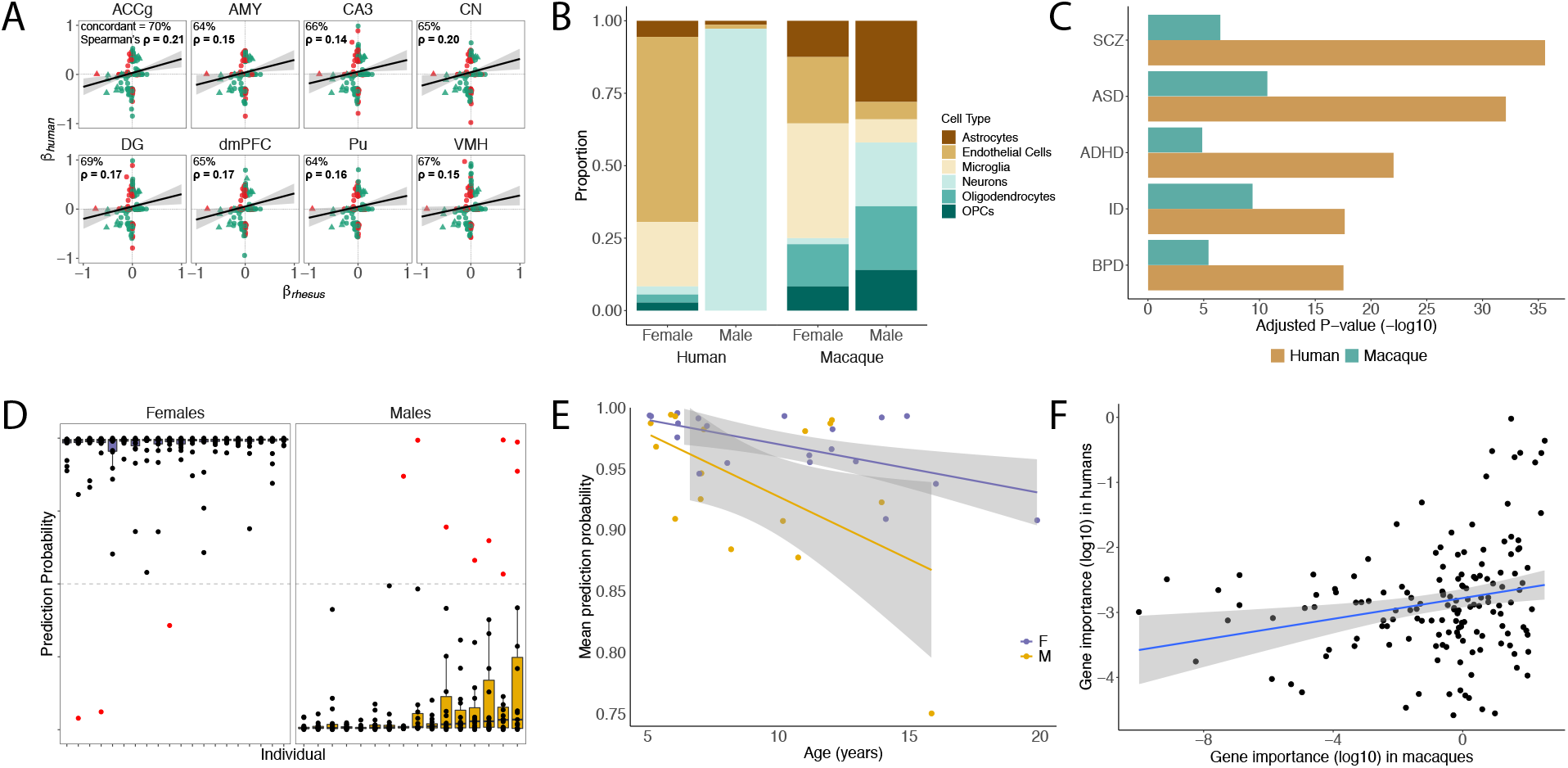
Sex-biased gene expression in macaque and human brains is associated with specific cell types and sex biased diseases. (**A**) Scatterplots of estimated sex effects for genes that are significantly sex-biased (LFSR < 0.05) in either humans (GTEx) or rhesus macaques (this study). Both autosomal genes (circles) and X chromosome genes (triangles) are included. Green points represent genes with concordant sex-bias across species, while red points represent discordance. Significant correlations (p < 0.05) are in bold. (**B**) Proportions of female-biased or male-biased genes that are also marker genes for specific cell types (see legend). Humans: female-biased genes (N = 36) are enriched for microglia (padj = 0.003) and endothelial cells (p_adj_ < 0.001) and male-biased genes (N = 73) are enriched for neurons (p_adj_ < 0.001). Macaques: female-biased genes (N = 47) are enriched for microglia (p_adj_ = 0.010) and male-biased genes (N = 51) are not enriched for any cell types. (**C**) Bar charts depicting adjusted p-values (-log10) from KS tests of male-biased disease risk gene enrichment for humans and macaques. SCZ = schizophrenia, ASD = autism spectrum disorders, ADHD = attention deficit hyperactivity disorder, ID = intellectual disability, BPD = bipolar disorder. (**D**) Boxplots of prediction probabilities of the known sex per individual (from models of all non-Y chromosome genes). Dots indicate values for individual samples. Purple boxes = female, yellow boxes = male, black dots = correctly classified samples, red dots = incorrectly classified sample (prediction probability of correct sex < 0.5). Boxplots indicate the median (black horizontal line), first and third quartiles (i.e., interquartile range, IQR; lower and upper hinges), and ranges extending from each to 1.5 x IQR beyond each hinge (whiskers). (**E**) Prediction probability (averaged across regions) of known sex per individual as a function of age (years) for females (purple) and males (yellow) from models of autosomal genes only (p (all) = 0.016). (**F**) Relative importance of X chromosome genes for sex prediction in X chromosome gene models (summed across regions) in the current study and Oliva et al. (2020) (ρ= 0.222, p = 0.006).

### Sex-biased brain gene expression partially reflects sex differences in microglial proportions

Observed sex differences in gene expression could reflect differences in cell composition and/or the expression of specific genes within cell types. To test the contribution of sex differences in cell type proportions to sex-biased gene expression in the macaque brain transcriptome, we drew on a recent meta-analysis of human brain cell type markers (41). We found that female-biased genes (LFSR < 0.05 in any region; n=270) were enriched for microglial marker genes (odds ratio [OR] = 2.51; p_adj_ = 0.003; Figure 3; Supplementary Table 7). Similarly, the largest sex differences in estimated cell type proportions (surrogate proportion variables, SPVs) were for microglia, with females exhibiting higher mean SPVs across regions (microglia: p_adj_ = 0.003; all other cell types: p_adj_ > 0.05) (Supplementary Figure 6, Supplementary Table 8). Female-biased genes identified in our analysis of the human GTEx data (Methods) were also enriched for microglial markers (OR = 4.607, p_adj_ = 0.003) (Figure 3; Supplementary Table 7). These results are consistent with previous reports of higher microglial proportions in the neocortices of adult female humans (42) and in multiple brain areas of adult female rats (43), in addition to a neurodevelopmental gene co-expression module in the human brain (ME3) that is enriched for both microglial markers and female-biased genes in the postnatal period (44). Differences in microglial number and maturation during development not only drive sexual differentiation in the brain, but are also likely to contribute to sex differences in AD and ASD (45, 46). Accordingly, we also estimated sex effects after performing cell type deconvolution analysis on the expression data (Methods; Supplementary Table 9). This analysis allowed us to identify sex-biased gene expression patterns that are not driven by sex differences in cell type abundances. Estimated sex effects tended to be in the same direction (i.e., male- or female-biased) whether or not cell type proportions were considered (74% concordance across all estimated effects; 99% concordance across effects that are significant in at least one analysis; ρ = 0.635, p < 2.20e-16; Supplementary Figure 7). As expected, fewer sex-biased genes pass our LFSR threshold in this analysis (25% fewer; N = 422 genes exhibited sex bias in at least one region) (Figure 2; Supplementary Results), 59% of which were also detected in at least one region in our primary analysis (Figure 2). Accordingly, we repeated all enrichment analyses described below using cell-type corrected data. The results were largely similar (Supplementary Results; Supplementary Figures 8-9; Supplementary Tables 16-18), so below we report only the results from our primary analyses.

### Sex-biased genes are involved in metabolic and immune-related pathways

Genes that exhibited female-biased expression in the macaque brain (LFSR < 0.05 in any region) were involved in pathways related to hormone receptor signaling, lipid catabolism, protein localization, translation, gliogenesis, apoptosis, inflammation (e.g., NF-κB signaling), and the immune response (e.g., monocyte differentiation, B cell production) (Fisher’s exact tests for these GO terms: p < 0.05) (Supplementary Table 10). These findings parallel those found in humans, where many genes that are more highly expressed in females are involved in immunity (14, 16, 47). This conserved pattern is likely to reflect evolved sex differences in immune surveillance and response due to the need for mothers to tolerate an internal, immunologically challenging pregnancy (48). Genes that exhibited male-biased expression in the macaque brain (LFSR < 0.05 in any region) were associated with pathways involved in lipid biosynthesis, vesicular transport, RNA localization, microtubule-based processes, immune effector processes, and cell cycle regulation (Supplementary Table 10). Sex differences in the expression of genes associated with the cell cycle and metabolism are consistent with findings in humans (14, 15, 47) and may reflect sex differences in brain size (49). The only pathway that was enriched in both male- and female-biased (non-overlapping) gene sets in macaques involves multivesicular bodies (underlying genes: male-biased = *TSPAN6, DENND10, VPS4B, TMEM50A*; female-biased = *CHMP4B, LAPTM4B*), which are critical for sorting and degradation of cellular proteins and play a role in neurodegenerative diseases, including AD (50).

### Sex-biased genes are implicated in sex-biased neurological disorders

Given that sex-biased genes in the human brain are linked to sex-biased neurological conditions (11–19), we investigated whether genes exhibiting normative sex differences in expression in the rhesus macaque brain also showed similar disease associations. Indeed, male-biased gene expression was linked to risk genes for ASD (KS tests on mean standardized β across regions: D = 0.148, p_adj_ = 2.14e-08), intellectual disability (D = 0.107, p_adj_ = 2.54e-07), schizophrenia (D = 0.098, p_adj_ = 1.31e-04), bipolar disorder (D = 0.162, p_adj_ = 9.15e-04), and ADHD (D = 0.146, p_adj_ = 0.003) (Figure 3; Supplementary Table 11). Within regions, these associations were the strongest in the anterior cingulate, dorsomedial prefrontal, primary visual, and superior temporal cortices, as well as the dentate gyrus (Supplementary Table 12). The cingulate cortex, superior temporal cortex, and hippocampus are critical for social valuation and interaction, and are among the most disrupted brain areas in schizophrenia (51). In addition, male-biased genes were associated with genes linked to AD in the anterior cingulate cortex only (Supplementary Table 12), a region that is commonly damaged across all neuropsychiatric AD symptoms (52). Although some male-biased genes are shared across all these conditions, many genes are unique to individual conditions or combinations of conditions (Supplementary Figure 10). Female-biased gene expression was not associated with any neurological conditions (Supplementary Table 11). These results are similar to studies of human brains, which have found that male-biased genes are associated with schizophrenia, bipolar disorder, AD, and ASD risk genes (15, 14, 16) (c.f. 11) and suggest that sex differences present in typically developing individuals modulate the impact of risk variants and contribute to the sex differences in disease prevalence. Consistent with this, we also found that male-biased gene expression in the human GTEx data was associated with risk genes for schizophrenia, bipolar disorder, ASD, ADHD, and intellectual disability (Figure 3; Supplementary Table 13). These results suggest that greater male susceptibility to certain conditions may be linked to male-biased expression of the risk genes for those conditions, and as a result, disruptive mutations in these genes have a larger impact in males (11).

To further investigate links between sex differences in brain gene expression and human disorders, we tested whether sex-biased genes in the macaque brain (LFSR < 0.05 in any region) were enriched for genes that exhibit altered expression levels in the brains of people with ASD (rather than ASD-risk genes identified in GWAS or twin studies). Female-biased genes were associated with cortex-wide ASD-upregulated genes that were recently identified in the largest ASD case-control study to date (53) (OR = 2.090, p = 0.0002), while male-biased genes were associated with ASD-downregulated genes (OR = 1.337, p = 0.035). This was also the case for sex-biased genes identified in our analysis of human GTEx data, which as expected, showed even stronger enrichments (female-biased and ASD-upregulated: OR = 14.906, p = 4.9e-21; male-biased and ASD-downregulated: OR = 4.719, p = 5.4e-15). These results are consistent with converging evidence of complex interactions between ASD, sex, microglia, and the immune system. In particular, previous studies have linked ASD-upregulated genes to microglial markers and immune genes (53–57) and suggest higher microglial proportions in ASD patients (53). These findings also parallel female-biased characteristics of macaque and human brains reported here and elsewhere (Supplementary Table 7) (14, 16, 42–44, 47), suggesting that ASD and female brains independently exhibit greater activation of the neuroimmune system. In fact, many female-biased, ASD-upregulated genes in both species were also microglial markers, and among genes identified as female-biased in either species (N = 249), genes that were also upregulated in ASD (N = 66) were enriched for numerous cytokine- and immune-related pathways (p < 0.05) (Supplementary Table 14). This bolsters previous work on rhesus macaque models of maternal immune activation-associated neurodevelopmental disorders (which include ASD) (58, 59).

Although these results appear to contrast with previous work linking male-biased genes in the typically developing human brain to microglial markers and ASD-upregulated genes (11, 60), differences in sample size and developmental period may explain the apparent discrepancy. For example, although our analyses include relatively more samples for both the ASD and normative sex datasets, these data represent different developmental periods (the ASD expression dataset analyzed here includes children and adults (53); the macaque and human GTEx datasets include adults only). This may impact results since ASD and sex both represent “developmentally moving targets” (60). Finally, more recent analyses (incorporating data from more individuals) report that ASD-upregulated microglial/immune gene modules are male-biased prenatally but are then female-biased during certain postnatal periods (60). This is consistent with prenatal male-biased and postnatal female-biased expression of a microglia-enriched neurodevelopmental gene module in the human brain (ME3) (44), in addition to reversal of sex differences in microglial colonization and activation prior to adolescence in rats (43). This suggests that ASD-upregulated microglial/immune genes that are expressed at higher levels in the adult female brain may be more highly expressed in the prenatal male brain during critical periods of neurodevelopment. These results may be interpreted as support for the gender incoherence (GI) theory (61) rather than the extreme male brain (EMB) theory (62) of ASD: while the EMB theory is consistent with reports of “masculinized” traits in individuals with ASD, including hyper-developed systemizing skills (62) (c.f. 63) and brain region-specific measures (64), the GI theory is supported by reports of “androgenized” traits in individuals with ASD, including digit ratios (61) and brain region-specific measures (64–66).

### Many sex-biased genes are regulated by estrogens

To illuminate the regulatory mechanisms underlying sex-biased gene expression in the macaque brain and to compare these mechanisms with those in humans, we identified motifs that tended to exist within the promoters of sex-biased genes more often than those of non-biased genes (Methods). We found that the promoters of sex-biased genes in the macaque brain are significantly enriched for estrogen receptor binding site motifs (Erra: OR = 1.066, p = 0.021) (Supplementary Table 15; Supplementary Figure 11). In further support of the importance of ER dynamics, we found that the most highly enriched motifs were for transcription factors that interact with estrogens, such as SF1 (OR = 1.349, p = 0.003) and Tbet (OR = 1.207, p = 0.003).

This is consistent with previous findings that, in humans, many sex-biased autosomal genes are indirectly modulated by sex hormones (14). Although previous work on sex differences in the human brain transcriptome associated these differences with androgen regulation (16), this study focused on sex-biased splicing patterns and included a large proportion of post-menopausal women (16). Finally, many other enriched motifs identified here also regulate sex-biased gene expression across human tissues (e.g., estrogens, *HNF4a, NRF1, IRF3, FOXA1*) (12), which is likely to reflect conserved regulatory mechanisms across species and tissues.

### Many sex-biased genes are regulated by estrogens

In order to investigate heterogeneity in sex-biased gene expression across individuals (of the same sex), and to identify potential drivers of this variation, we constructed and evaluated region-specific sex prediction models of the rhesus macaque brain transcriptome (model construction repeated using 3 gene sets [non-Y chromosome genes (Figure 3), X chromosome genes only, autosomal genes only] within each region, resulting in 15 regions x 3 gene sets = 45 models total). We could accurately predict sex from the expression levels of relatively few genes (models of non-Y chromosome genes: mean accuracy = 0.977, mean number of genes = 39; models of X chromosome genes only and autosomal genes only performed similarly; Supplementary Table 19), similar to previous work across human tissues (12). While most genes (87%) were only influential in one region, they tended to be ubiquitously expressed (89% expressed in at least 13 regions), which is likely to reflect that the magnitude of sex differences in expression per gene varies across brain regions, even for shared sexbiased genes (Supplementary Table 20). These models (all non-Y chromosome) tended to be better at correctly classifying female individuals (Figure 3; Supplementary Table 19; Supplementary Figure 12), which may reflect that, of genes that were influential in at least one region (N = 501), X chromosome genes were more influential than autosomal genes (average of summed relative influence: X chromosome: mean = 14.28; autosomes: mean = 8.42) (Supplementary Tables 19-20). Accuracy was also lower in predicting the sex of older individuals (linear regression of known sex probability modelled as a function of age: p = 0.016) (Supplementary Figure 13), specifically in models of autosomal genes (Figure 3). This effect was stronger among males of all ages and older individuals (>8 years) (Supplementary Figures 13, 14). In fact, the most often misclassified individual was also the oldest male in our sample (misclassified as female in 7/45 models, spanning 3 different regions and all gene sets; out of 527 samples x 3 gene sets = 1581 classifications, there were only 16 misclassifications total) (Supplementary Table 21). These sex and age differences in prediction accuracy may reflect that males, particularly older males (>8 years), exhibit higher within-sex gene expression variation compared to females (median pairwise Euclidean distance of residual gene expression among: old males = 137.9, young males = 135.9, old females = 134.2, young females = 133.7; all differences are significant except young males versus old females, Tukey’s HSD p_adj_ < 0.05; Supplementary Figure 15).

Models of X chromosome genes highlighted similarities with humans, as that the most influential genes in this study were also the most influential genes in models constructed in a recent study across 44 human tissues (12) (N = 150 one-to-one orthologues with nonzero influence in both studies; ρ = 0.222, p = 0.006; Figure 3; Supplementary Figure 16). This reflects species similarities in the magnitude of sex-biased expression across X chromosome genes (ρ > 0.69 across 8 overlapping regions with the human GTEx data; Figure 3; Supplementary Figure 5; Supplementary Table 6). Female samples that were misclassified in X chromosome gene models tend to exhibit relatively low expression of the most influential X chromosome genes in those regions (Supplementary Figure 17), which may partially reflect variability in XCI escape across female individuals and tissues (67).

### Many sex-biased genes are regulated by estrogens

To better understand the evolutionary dynamics underlying sex differences in the macaque brain transcriptome, we examined four mechanisms that may facilitate the rapid evolution of sex-biased gene expression observed in other studies: i) their tendency to be located on sex chromosomes (due to sex-specific patterns of selection and inheritance (68)), since the smaller effective population size of these chromosomes may lead their genes to evolve more rapidly (69); ii) higher tissue specificity (i.e., lower pleiotropy) (70, 71) since pleiotropy may constrain evolutionary change due to widespread multivariate stabilizing selection (72, 73); iii) higher genetic variance in gene expression, since genes whose expression is attributable to genetic variance (vs. environmental variance) can better respond to selection (71); and iv) higher genic tolerance, since this would allow for more coding sequence mutations without losing function.

We found that sex-biased genes tend to be located on the sex chromosomes (mechanism i above) and that sex differences in expression predict tissue specificity (ii) and genetic variance (iii), but not genic tolerance (iv). Specifically: i) the X and Y chromosomes were enriched for sex-biased genes (X chromosome: OR = 2.16; p_adj_ = 0.002; Y chromosome: OR = Inf; p_adj_ < 0.001), and these enrichments were driven by female- and male-biased genes, respectively (X chromosome female-biased: OR = 2.79; p_adj_ = 0.004; X chromosome male-biased: OR = 1.62; p_adj_ = 1; Y chromosome expression is male-specific) (Figure 2). Female-biased gene enrichment on the X chromosome is consistent with a preponderance of female-beneficial mutations that are dominant, since these mutations occur in females two-thirds of the time and are, therefore, selected for (in females) more often than selected against (in males) (68); ii) Tissue specificity estimates ranged from 0.018 to 1 (mean = 0.172; sd = 0.148; Supplementary Table 22) and genes exhibiting larger sex differences in residual expression also showed more tissuespecific expression (ρ = 0.332; p < 2.2e-16) (Figure 4); iii) The structure of our data resulted in a bimodal distribution for estimates of genetic variance (*vu*), so we evaluated the relationship between log(*vu*) and sex differences in residual expression separately within each distribution, and found significant positive associations in both (upper distribution: ρ = 0.234, p < 2.2e-16; lower distribution: ρ = 0.290, p < 2.2e-16) (Figure 4); and iv) We did not detect a relationship between absolute sex differences in residual expression and LOF mutation tolerance (ρ= 0.006, p = 0.622) (Figure 4).

**Figure 4.**
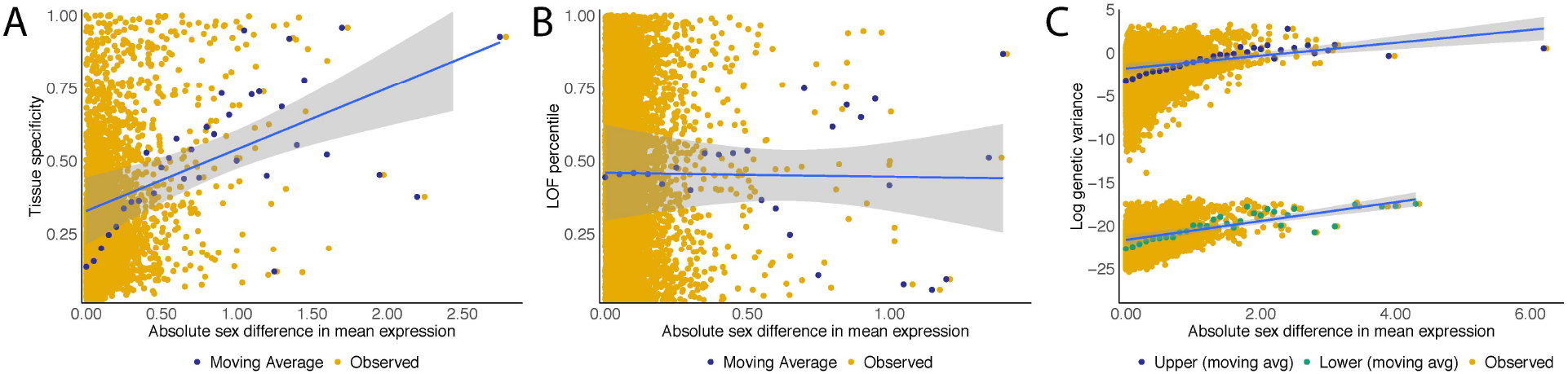
Evolutionary characteristics of sex differences in gene expression. (**A**) Tissue specificity as a function of the absolute difference in mean residual expression per gene (averaged across regions) (N = 12,663, excludes Y chromosome genes). (**B**) Loss-of-function (LOF) tolerance as a function of the absolute difference in mean residual expression per gene (averaged across regions) (N = 7,786, includes one-to-one orthologues in LOFtools database only, excludes Y chromosome genes). (**C**) Genetic variance (log) as a function of the absolute difference in mean residual expression per gene and region (N = 152,431 excluding Y chromosome genes).

## Conclusions

This work provides an in-depth characterization of the patterns, biological functions, disease associations, regulatory factors, and evolutionary mechanisms relevant to sex-biased gene expression in the rhesus macaque brain. We highlight that sex-biased genes exhibit greater genetic variance for expression, more tissue-specific patterns of expression, and a tendency to be located on sex chromosomes. Despite the presence of these factors, which are likely to drive evolutionary divergences in the expression and function of sex-biased genes across species, we found that humans and rhesus macaque brains exhibit similar transcriptomic sex differences. Not only are gene expression levels biased in the same direction (i.e., female- or male-biased) in multiple brain areas, but adult macaque and human brains appear to share estrogen-mediated regulation of sex-biased genes, upregulation of the neuroimmune system in females, and sex-biased expression of genes implicated in sex-biased conditions, including ASD. These similarities bolster the translatability of this indispensable model species for studies of sex-biased neurological conditions. The dataset generated here represents a valuable resource for future studies of the rhesus macaque brain transcriptome, and future work on earlier developmental periods will further elucidate the extent to which sex differences in the expression of human disease-linked genes are conserved across species.

## Acknowledgements

We thank those who make our research possible, particularly the Caribbean Primate Research Center and staff from both the Cayo Santiago Field Station and Sabana Seca Field Station. We also thank Madeleine Rosenstein and for assisting in data collection and Dr. Armin Raznahan for providing valuable feedback.

The Genotype-Tissue Expression (GTEx) Project was supported by the Common Fund of the Office of the Director of the National Institutes of Health (commonfund.nih.gov/GTEx). Additional funds were provided by the NCI, NHGRI, NHLBI, NIDA, NIMH, and NINDS. Donors were enrolled at Biospecimen Source Sites funded by NCI\Leidos Biomedical Research, Inc. subcontracts to the National Disease Research Interchange (10XS170), Roswell Park Cancer Institute (10XS171), and Science Care, Inc. (X10S172). The Laboratory, Data Analysis, and Coordinating Center (LDACC) was funded through a contract (HHSN268201000029C) to The Broad Institute, Inc. Biorepository operations were funded through a Leidos Biomedical Research, Inc. subcontract to Van Andel Research Institute (10ST1035). Additional data repository and project management were provided by Leidos Biomedical Research, Inc.(HHSN261200800001E). The Brain Bank was supported by supplements to University of Miami grant DA006227. Statistical Methods development grants were made to the University of Geneva (MH090941 & MH101814), the University of Chicago (MH090951,MH090937, MH101825, & MH101820), the University of North Carolina Chapel Hill (MH090936), North Carolina State University (MH101819),Harvard University (MH090948), Stanford University (MH101782), Washington University (MH101810), and to the University of Pennsylvania (MH101822). The datasets used for the analyses described in this manuscript were obtained from dbGaP at http://www.ncbi.nlm.nih.gov/gap through dbGaP accession number phs000424.v8.p2.

## Competing interests

The authors have no competing interests to declare.

Data and materials availability: All code used and data generated in this study will be made available upon publication.

## Author Contributions

ARD, JPH, CCS, KLC, and NSM conceptualized the research; MJM, NSM, KLC, OG, and SEBS and collected brain tissue, facilitated by MIM, AVRL, JS, CSW, SCA, MKS, JPH, MLP, and CBRU; MAW contributed data; ARD, KLC, and AM and performed genomic lab work; JEN collected behavioral data using a protocol designed by LJNB; ARD and KLC performed genomic analysis with input from NSM; CT and LJNB performed behavioral analysis; ARD, KLC, JPH, and NSM wrote the manuscript; All authors reviewed and revised the manuscript.

## Methods

### Sample collection

#### Tissue procurement and processing

Rhesus macaque (*Macaca mulatta*) individuals were from the Cayo Santiago population and were removed from the island and humanely euthanized as part of the population management strategy implemented by the Caribbean Primate Research Center, University of Puerto Rico. No individuals were used for brain invasive procedures or had signs of malformations or lesions. Within 30 minutes of euthanasia, following perfusion with cold saline, whole brains were extracted. Left and right hemispheres were separated using a sterilized razor, and left hemispheres were set aside for fixation. Right hemispheres were placed in a mold and cut into ½ centimeter coronal slabs. Slabs were flash frozen using an ethanol and dry ice mixture. Brains were stored at -80°C until dissection.

#### Tissue dissection

Brain samples were collected postmortem from 36 adult macaques (20 females, 16 males). Frozen slabs were kept on dry ice during sampling. Samples were collected using 1mm surgical punches with reference to coronal cross-sections from the rhesus macaque anatomical brain atlas (74). This sampling method allowed us to sample relatively evenly across all cortical layers, which exhibit distinct cell composition and gene expression patterns (75).

Fifteen brain regions of interest were identified on frozen hemispheres using gross landmarks (e.g., cortical sulci/gyri and white matter tracts). Specifically: 1) The ventromedial prefrontal cortex (vmPFC; areas 10m/32) was sampled from the rostral most plane when the cingulate, principal, and medio-orbital sulci were visible. Six punches were taken on the cingulate gyrus, from slightly inferior to the tip at the medial surface towards the underlying white matter in the latero-inferior direction; 2) The dlPFC (area 46d) was sampled from the rostral most plane when the cingulate, principal, and medio-orbital sulci were visible. Six punches were taken on gyrus superior to the principal sulcus, from the tip at the lateral surface towards the underlying white matter in the medio-inferior direction; 3) The vlPFC (area 12r) was samples from the rostral most plane when the cingulate, principal, and medio-orbital sulci were visible. Six punches were taken on gyrus inferior to the principal sulcus, from the tip at the lateral surface towards the underlying white matter in the medial direction; 4) The dmPFC (area 9m) was sampled from rostral most plane when the cingulate, principal, and medio-orbital sulci were visible. Six punches were taken on the superior frontal gyrus, from the tip at the medial surface towards the underlying white matter in the latero-inferior direction; 5) The anterior cingulate gyrus (ACCg; area 24) was sampled from the rostral most plane when the corpus callosum was visible. Six punches were taken on the cingulate gyrus, from the tip at the medial surface towards the underlying white matter in the latero-inferior direction; 6) The mid-superior temporal sulcus (mid-STS) was sampled when the central, intraparietal, lateral, and superior temporal sulci were visible. Six punches were taken from the inferior-most point of STS towards the superior white matter (Sallet et al. 2011); 7) The primary motor cortex (M1; area 4) was sampled when the precentral, central, lateral, and superior temporal sulci were visible. Six punches were taken from the superior-medial portion; 8) The primary visual cortex (V1; area 17) was sampled on the anterior surface of the most posterior slab, in the inferior arm of the calcarine sulcus; 9) The caudate nucleus (CN) was sampled at the rostral most point at which the internal capsule was visible and clearly separated the caudate nucleus from the putamen. Six punches were taken from the most superior-lateral point moving in the inferomedial direction; 10) The putamen (Pu) was sampled at the rostral most point at which the internal capsule was visible and clearly separated the caudate nucleus from the putamen. Six punches were taken from the most superior-lateral point moving in the inferomedial direction; 11) The amygdala was sampled when both it and optic chiasm were clearly visible. Seven punches were taken lateral to optic chiasm, sampling across the superior portion of the amygdaloid complex (representing the anterior cortical nucleus, central nucleus, medial nucleus, and superior portions of the accessory basal and basal nuclei); 12) The dentate gyrus (DG) was identified within the hippocampal formation by its slightly darker color, caused by a high density of small granule cells. This sampling also included CA4, which is difficult to differentiate from the polymorphic layer of the DG. Accordingly, other studies have combined these regions, collectively referring to them as the hilus (which is the formal name for the polymorphic layer of the DG) (76); 13) The CA3 was sampled from the area superior to the DG within the hippocampal formation, in the mediolateral direction. This sampling likely also included portions of CA2 and CA1; 14) The lateral geniculate nucleus of the thalamus (LGN) was sampled when it was clearly visible, and six punches were taken across all layers in the medio-inferior direction; 15) The ventromedial hypothalamus (VMH) was sampled from rostral most plane when the hypothalamus was visible. This sampling likely also included portions of surrounding nuclei (e.g., arcuate). Four to six punches were taken from the most medio-inferior portion. Dissected tissue samples were stored at -80ºC prior to further processing.

### Bulk RNA-seq data generation and quality assessment

#### RNA extraction

1 mL of Trizol was added to dissected frozen tissue samples immediately before lysing. A single chilled 5mm stainless steel bead was added to each tube before placing samples in the TissueLyser II bead mill. Samples were homogenized for 2 minutes at 20 hz. Plates were rotated before homogenization was repeated. Homogenized samples were then transferred to new tubes and incubated at room temperature for 5 minutes. 200 mL of chloroform was added to each sample, tubes were manually shaken for 15 seconds, incubated at room temperature for 2 minutes, and centrifuged at 12k at 4ºC for 15 minutes. The upper aqueous solution (containing RNA) was transferred to a new sample tube, and Total RNA was extracted using Zymo Quick-RNA Microprep kits. Each sample was subjected to DNase treatment as per manufacturer’s instructions.

#### RNA quality assessment

RNA quality was assessed using a Fragment Analyzer or a Tapestation, which provided RQN or RINe values, respectively. For later analyses, RQN and RINe values were converted to RIN values using published regression lines (RQN = 0.9697*RIN, R2 = 0.9635; RINe = 0.991*RIN, R2 = 0.936) (77, 78).

#### Library preparation and sequencing

cDNA libraries were prepared using the NEBNext Ultra II RNA Library Prep Kit for Illumina, as per the manufacturer’s instructions with some modifications. Briefly, poly-adenylated mRNA was purified from 200 ng of total RNA using the NEBNext Poly(A) mRNA Magnetic Isolation Module. The mRNA was then reverse transcribed into cDNA, ligated to Illumina adapters, size-selected for a median size of 600 bp, and amplified via PCR for 12 cycles. Each sample was tagged with a unique molecular barcode and pooled samples into Illumina NovaSeq lanes (across 2 sequencing runs, one using 2×50bp sequencing on the S2 flow cell and another using 2×100bp sequencing on the S4 flow cell).

#### Reference genomes and read alignment

Following sequencing, we mapped reads to the rhesus macaque transcriptome v10 (Ensembl) using the pseudoaligner kallisto v0.43.1 (79). Given that sequence homology across the sex chromosomes present in reference genomes/transcriptome can lead to technical mapping errors, we created two modified, sex-specific transcriptomes and separately mapped reads from males and females (following 80). Specifically, the Y chromosome was removed from the female-specific transcriptome, and *CD99* on the Y chromosome (within the pseudoautosomal region (81)) was removed from the male-specific transcriptome. We also confirmed chromosomal sex of individuals by mapping to a non-sex-specific transcriptome and examining Y chromosome gene counts.

We imported the transcript count matrices for males and females into R using the function tximport (R package tximport) and combined them into one count matrix. We summarized transcript counts to the gene level using the appropriate functions in the R package biomaRt and the function summarizeToGene (R package tximport). This procedure resulted in a 22,514 × 532 (p x n) read-count matrix, where p is the number of genes measured and n is the number of samples. We confirmed the identity of all samples based on genotyping from the RNA-seq reads.

#### Quality assessment

We removed 5 samples that were low quality (e.g., samples with low Phred scores and/or high PCR duplication rates). This resulted in a 22,514 × 527 (p x n) read-count matrix, where p is the number of genes measured and n is the number of samples. We also confirmed the chromosomal sex of all individuals/samples by mapping to an unedited (non-sex-specific transcriptome) and examining Y chromosome gene expression (Supplementary Figure 18).

#### Read normalization

We normalized the read count matrix using the functions calcNormFactors (R package edgeR (82) and voom in the R package limma (83). Prior to further RNA-seq data analysis, we filtered out genes that were very lowly or not detectably expressed in our samples. Specifically, within each region we removed any gene with mean TPM<10 in both males and females (i.e., genes with >=10 mean TPM in at least one sex were retained). This procedure resulted in a mean of 10,171 genes (range: 9,617-11,135), and 12,672 unique genes were detectably expressed in at least one brain region. These data (normalized log2 counts per million reads) were used throughout the statistical analyses described below.

#### Genotyping

We used genotype data (with variants called from RNAseq data) to control for genetic relatedness among individuals in this study. For each sample, we mapped reads to the rhesus macaque reference genome v10 (Ensembl) using STAR (84) and then pooled mapped reads for each individual across all brain regions. We used the Genome Analysis Toolkit (GATK) (53, 54) to mark duplicates (MarkDuplicates), split reads spanning splice events (SplitNCigarReads), and recalibrate base quality scores (BaseRecalibrator and ApplyBQSR) before calling variants (HaplotypeCaller) using a standard minimum confidence threshold for calling of 20.0. We retained sites that passed the following filters: QD < 2.0; MQ < 40.0; FS > 60.0; HaplotypeScore > 13.0; MQRankSum < -12.5; and ReadPosRankSum < -8.0. We estimated kinship with the program lcMLkin using variants that were genotyped in all 36 individuals, had minor allele frequencies > 0.3, minimum completeness of 0.9, and were at least 100 kb apart (thinned using VCFtools (55)). These relatedness estimates were confirmed using known mother-offspring pairs (5 known pairs: mean relatedness estimate = 0.48; remaining pairs: relatedness estimates <= 0.25).

### Behavioral data collection

Previous work has shown that dominance rank can impact gene expression in the brain and peripheral tissues of wild and laboratory animals (e.g., 85–87). Here, dominance rank reflects the direction and outcome of win-loss agonistic interactions (e.g., aggression, threats, displacements, submissions) recorded during focal animal samples or during ad libitum observations. To calculate individual dominance ranks, behavioral data were collected for all animals in this study (and all other members of this social group age 4 and above) in the three months prior to removal. Methods for behavioral data collection as well as dominance rank inference in this population are described by Testard and colleagues (89, 90). Ranks were calculated separately within each sex because dominance is attained differently in male and female macaques. Specifically, male macaques tend to disperse from their natal groups and their rank in the new groups are largely determined by their duration of tenure (89, 90). Female macaques are philopatric and dominance rank is inherited maternally, resulting in stable linear dominance hierarchies among females (89, 90). Accordingly, known maternal relatedness was used to resolve behavioral gaps in the female hierarchy. To account for group size, dominance rank was first defined as the percentage of same sex individuals that a subject outranked. We then followed previous work (88) in creating categorical dominance ranks, calculated by classifying animals as high- (rank >= 80%), mid- (50% <= rank < 80%), or low-ranking (rank < 50%) based on their percentage dominance ranks within each sex. We modeled categorical dominance rank as an ordinal variable for all differential expression analyses using the ordered factor class in R.

### Statistical analyses

All statistical analyses were performed using R v4.0.0 (91) or Homer v4.10 (92).

#### Dimensionality reduction

To visualize the structure of the expression data, we applied dimension reduction methods to the normalized, filtered expression matrix. Prior to dimension reduction, the effects of library batch and RIN were removed from the data using the removeBatchEffect function in the R package limma (83). Dimension reduction was performed using Uniform Manifold Approximation and Projection (UMAP) via the umap function in the R package umap (93) with the following metrics: n_neighbors = 200, min_dist = 0.5, metric = ‘manhattan’. We also provide t-SNE and PCA plots in the supplement using the Rtsne function (perplexity=30) in the R package Rtsne (94) and the prcomp function in the R package stats).

#### Hierarchical clustering

Unsupervised hierarchical clustering was conducted using the normalized, filtered expression matrix. Prior to hierarchical clustering, the effects of library batch and RIN were removed from the data using the removeBatchEffect function in the R package limma (83). Cluster analyses were performed by the pvclust function (R package pvclust) (95). Correlation was used as the distance measure. This function provides both approximately unbiased (AU) p-value and bootstrap probability (BP) value. AU values are calculated using multiscale bootstrap resampling, while BP values are calculated by the ordinary bootstrap resampling (95). This method was applied to expression values averaged across samples per region (to examine clustering by region).

#### Variance partitioning

We performed variance partitioning on the normalized, filtered expression matrix using the fitExtractVarPartModel and plotVarPart functions in the R package variancePartition (96). This function allowed us to fit a linear mixed model to estimate contribution of multiple sources of variation while simultaneously correcting for all other variables. Prior to hierarchical clustering, the effects of library batch and RIN were removed from the data using the removeBatchEffect function R package limma (83). We modelled the expression of each gene as a function of individual, region, sex, age, and ordinal rank. Categorical terms were modeled as random effects, as recommended by the package’s creator (97). We then extracted and visualized the fraction of variance explained by each biological or demographic term, in addition to the residual variance.

#### Modeling sex effects on gene expression

To identify genes that were affected by sex within each region, we used linear mixed effects models that control for relatedness. We analyzed each of the 15 brain regions separately using the emmreml function in the R package EMMREML (37). Normalized gene expression values were modelled as a function of sex, age, ordinal rank, RIN, and library batch. Although standard normalizations fail to account for the effects of RNA degradation, statistically controlling for RNA quality corrects for most of these effects (98). For each gene in the normalized, filtered expression matrix, we estimated the effect of sex on gene expression using the Equation 1 below:

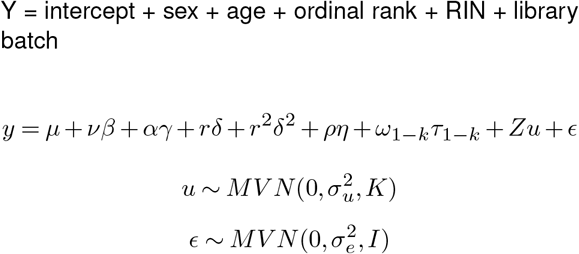

where y is the n by 1 vector of normalized gene expression levels for the n samples collected per region; μis the intercept; νan n by 1 vector of sex and βis its effect size; αis an n by 1 vector of age in years at the time of sample collection and γis its effect size; r is an n by 1 vector of linear contrasts of sex-specific rank and δis its effect size; r^2^ is an n by 1 vector of quadratic contrasts of sex-specific rank and δ2is its effect size; ρis an n by 1 vector of RIN values and ηis its effect size; and ω_1-k_ are k vectors (with k equal to the number of library batches for the given region), each of which is an n by 1 vector of a dummy variable for that library batch (0 = sample not included in this batch; 1 = sample included in this batch), and τ_1-k_ are the effect sizes for each vector. The m by 1 vector u is a random effects term to control for kinship and other sources of genetic structure. Here, m is the number of unique individuals sampled for each region, the m by m matrix K contains estimates of pairwise a relatedness derived from a genotype data set, σ_u_^2^ is the genetic variance component (0 for a non-heritable trait), and Z is an incidence matrix of 1’s and 0’s that maps samples to individuals in the random effects term. Residual errors are represented by ε, an n by 1 vector, where σ_e_^2^ represents the environmental variance component (unstructured by genetic relatedness), I is the identity matrix, and MVN denotes the multivariate normal distribution.

#### Multivariate adaptive shrinkage (MASH)

To identify genes that are differentially expressed between males and females and whether or not these effects are shared or region-specific sex effects, we used the outputs from the EMMA mixed models described above (i.e., per gene betas and their standard errors within each of 15 regions) as inputs for multivariate adaptive shrinkage models (R package mashr) (38). For missing data, betas were set to 0 and standard errors were set to 100 (as recommended by the mashr package’s creators). We first selected strong signals by running a condition-by-condition (1by1) analysis on all the data (mash_1by1 function) and extracting those results with local false sign rate (LFSR) < 0.05 in any condition. Specifically, this analysis runs ash in the R package ashr (99) on the data from each condition, an Empirical Bayes approach to FDR analysis that incorporates effect size estimates and standard errors, and assumes the distribution of the actual effects is unimodal, with a mode at 0 (39). We also generated a random subset of the data (50% of expressed genes), computed a list of canonical covariance matrices (cov_canonical function), and used these data and matrices to estimate the correlation structure in the null tests (estimate_null_correlation function). We then set up the main data objects (i.e., “strong” and “random”) with this correlation structure in place (mash_set_data function). We used the strong tests to set up data-driven covariances by performing PCA on the data (using 5 PCs; cov_pca function) and using the resulting 5 candidate covariance matrices to initialize and perform “extreme deconvolution” (cov_ed function) (100). We then estimated canonical covariances from the random tests and then fit mash to the random tests using both datadriven and canonical covariances. We extracted the fitted g mixture from this model and specified this mixture model when fitting mash to the strong tests. Significant genes (i.e., ‘sex-biased genes’) passed an LFSR cutoff of 0.05.

#### Human (GTEx) comparison

We estimated sex effects across 10 tissues from the human GTEx data (V8), including the amygdala, BA24, caudate nucleus, cerebellar hemisphere, BA9, hippocampus, hypothalamus, nucleus accumbens, putamen, and substantia nigra (mean N = 39F/119M; Supplementary Table 23). Technical replicates for two regions (“Cortex” and “Cerebellum”) were excluded. Using the EMMA models described above, we modelled gene expression (within each region and for each gene) as a function of sex, age, RIN, experimental batch, and ischemic time. We then applied MASH to the model outputs (as described above). To test for the consistency of sex effects on gene expression across data sets, we compared the results across 8 overlapping regions (AMY/amygdala, ACCg/BA24, CN/caudate, dmPFC/BA9, DG/hippocampus, CA3/hippocampus, VMH/hypothalamus, Pu/putamen) for all one-to-one orthologs that were significant (LFSR < 0.05) in at least one data set. We report Spearman’s rank order correlation coefficients (ρ) and the quadrant count ratio (q = (N concordant - N discordant) / total).

#### Cell type enrichment analysis

We tested for cell type enrichment among male- and female-biased genes using cell type markers from the R package BRETIGEA (BRain cEll Type specIfic Gene Expression Analysis) (41). In this package, the ‘markers_df_brain’ data frame contains the top 1000 marker genes (ranked by specificity) from each of the six major brain cell types (i.e., astrocytes, endothelial cells, microglia, neurons, oligodendrocytes, and OPCs), which were estimated from their meta-analysis of brain cell gene expression data from both humans (*Homo sapiens*) and mice (*Mus musculus*). Homo sapiens gene names were converted to (*Macaca mulatta*) Ensembl gene IDs using the bioMart R package. Sex-biased gene sets included any genes that were significantly male-biased or female-biased in any region (LFSR < 0.05). Fisher’s exact tests were used to test for cell type-specific enrichments (fisher.test function in the R package stats; alternative = ‘greater’). P-values were adjusted using the Benjamini-Hochberg method, and tests with adjusted p-values less than 0.05 were considered significant. This analysis was also performed on sex-biased genes (LFSR < 0.05 in any region) identified in our analysis of the human GTEx data (described above).

#### Deconvoluting cell type proportions and modelling sex effects on gene expression

Given that sex-biased gene sets were enriched for certain cell types (see Results), we also estimated sex effects after performing cell type deconvolution analysis in the R package BRETIGEA (41). Using the cell type marker genes described above, cell type deconvolution analyses were conducted within each region. First, the effects of library batch and RIN were removed from each normalized, filtered expression matrix using the removeBatchEffect function (R package limma). This matrix and the marker gene list were used as inputs to estimate the relative cell type proportions (i.e., surrogate cell type proportion variable (SPVs) for each cell type). This was performed by the findCells function, using the top 50 markers for each cell type and the singular value decomposition (SVD) dimension reduction approach, and scaling the gene expression data from each marker gene prior to using it as an input for dimension reduction. SPVs are eigenvectors of an SVD and do not directly quantify cell type proportions; rather, SPVs reflect relative differences in cell type composition within each cell type and, therefore, some SPVs will take on negative values. Finally, we adjusted each row of gene expression for sample differences in relative cell type proportions using the adjustCells function, which outputs the residuals from a linear model for downstream analysis. For each gene in the adjusted expression matrix, we estimated the effect of sex on expression using Equation 2 below (see ‘Modelling sex effects on gene expression’ section for details). Technical effects (i.e., library batch and RIN) were not included here since they were removed prior to the estimation of relative cell type proportions.

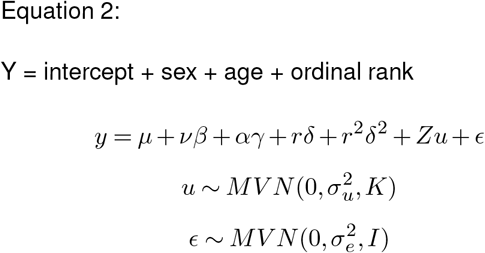

We then used the outputs from these models (i.e., per gene betas and their standard errors within each of 15 regions) as inputs for multivariate adaptive shrinkage models (see ‘Multivariate adaptive shrinkage’ section above for details). Significant genes (i.e., ‘sex-biased genes’) passed an LFSR cutoff of 0.05.

#### Functional enrichment analysis

Gene ontology (GO) enrichment analyses were performed using the R packages topGO (101) and ViSEAGO (102). GO term names were obtained from Ensembl using the Ensembl2GO and annotate functions. Enrichment analyses were conducted on male-biased and female-biased genes separately, with each set of genes defined as those that were significantly biased in mashr (LFSR < 0.05) in any region (excluding Y chromosome genes, which are not expressed in females). For each test, background genes represented all expressed genes that were not in the male- or female-biased gene set of interest. We used Fisher’s exact tests, which are based on gene counts. Enrichments with nominal p<0.05 were considered significant, as suggested by the topGO package’s creators. The parent child algorithm (103) was used since it determines over-representation of terms in the context of annotations to the term’s parents. Other approaches to measuring over-representation of GO terms cannot cope with the dependencies resulting from the structure of GO because they analyze each term in isolation. The parent child approach reduces the dependencies between the individual term’s measurements, and thereby avoids producing false-positive results owing to the inheritance problem. We computed the semantic similarity between GO terms using Wang’s method (compute_SS_distances function) (104), and clustered GO terms using Ward’s clustering criterion (GOterms_heatmap function) (105).

#### Disease gene set enrichment analysis (GSEA)

Gene set enrichment analyses for disease ontology (DO) terms were performed using human risk genes downloaded from the DISEASES resource, which integrates the results of text mining and manually curated disease-gene associations, cancer mutation data, and genome-wide association studies from existing databases (108). (*Macaca mulatta*) Ensembl IDs were linked to human diseases from this database using one-to-one human orthologues (and their associated proteins) from the R package bioMart. Diseases with at least 10 associated genes were retained for further analysis. Genes were ranked according to their mean standardized betas (i.e., mashr beta divided by posterior SD, averaged across regions). For each disease in the final dataset (N=1257), we used two-sample Kolmogorov-Smirnov (KS) tests to compare the cumulative distribution functions of the mean standardized betas of genes associated with the disease to those associated with every other disease in the data set. We tested two alternative hypotheses, namely that the cumulative distribution function for the target set was either less than or greater than that of the background set. P-values were adjusted using the Benjamini-Hochberg method, and tests with adjusted p-values less than 0.05 were considered significant. These analyses were also run within each region (using ranked standardized betas), and on the mean standardized betas from our analysis of the human GTEx data (described above).

We also tested whether the sex-biased genes identified here tend to exhibit altered expression levels in human disease. Whole cortex ASD-upregulated and -downregulated genes were collected from (53). (*Macaca mulatta*) Ensembl IDs were linked to human diseases from this database using one-to-one human orthologues (and their associated proteins) from the R package bioMart. For each ASD gene set, Fisher’s exact tests were performed on female- and male-biased gene sets. This analysis was also performed on the standardized sex effects estimated in our analysis of the human GTEx data (described above).

#### Motif enrichment analysis

We used HOMER (Hypergeometric Optimization of Motif EnRichment) (92) to analyze the promoters of genes and look for motifs that are enriched in the target gene promoters relative to other promoters. The target gene set consisted of all genes that were consistently sexbiased in at least 13 regions. We searched for motifs from -1000 to +300 relative to the transcriptional start site (TSS) using HOMER’s curated set of 414 known vertebrate motifs. The program assigns weights to the background promoters based on the distribution of GC content in the target gene promoters to ensure that comparable numbers of low and high-GC promoters are analyzed. It also performs autonormalization to remove sequence content bias from lower order oligos (1/2/3-mers) by adjusting background weights based on the target distribution. The hypergeometric distribution is used to score motifs. Enrichment p-values less than 0.05 were considered significant. QQ plots of observed versus expected -log10 p-values include inflation estimates (lambda = the median of the resulting chisquared test statistics divided by the expected median of the chi-squared distribution; <1 = less significant than expected; 1 = in line with uniform distribution; >1 = more significant than expected).

#### Sex prediction

Sex prediction: For each region, we created sex prediction models using residual gene expression values (i.e., expression levels after removing the effects of age, ordinal rank, RIN, library batch, and relatedness using the EMMA models and the normalized, filtered expression matrix described above). Specifically, we implemented gradient boosted models (GBM) using leave-one-out cross validation in the R package caret. We fit these models across various tuning parameters (interaction depths = 1, 3, 5, 9; number of trees = 50, 100, 150, 200, 250; shrinkage = 0.1; n.minobsinnode = 5), and models with the highest receiver operating characteristic (ROC) values were selected as the optimal model for each region. For each optimal model, we extracted the prediction probabilities for each sample, calculated the relative influence of each gene using the GBM model based technique in the varImp function (i.e., relative influence = the reduction in sums of squared error due to any split on that predictor, summed over all trees in the model (107)) scaled to a maximum value of 100) and calculated multiple performance metrics, including accuracy, sensitivity, and specificity (see below). This was done separately for: 1) combined X chromosome and autosomal genes; 2) autosomal genes only; and 3) X chromosome genes only.

True “Positives” (TP) = the number of samples correctly identified as female

True “Negatives” (TN) = the number of samples correctly identified as male

False “Positives” (FP) = the number of samples incorrectly identified as female

False “Negatives” (FN) = the number of samples incorrectly identified as male

Accuracy = (TP+TN) / (TP+TN+FP+FN)

Sensitivity = TP / (TP+FN)

Specificity = TN / (TN+FP)

We tested for age effects on the accuracy of our sex prediction models by modelling mean prediction probability of known sex per individual as a function of age. We also examined age effects on the variability of our sex predictions by modelling the standard deviation of known sex prediction probabilities per individual as a function of age. To test for the stability of these effects within old and young individuals and for varying sample sizes, we examined age effects on accuracy within subsamples of young (<= 8 year) and old (>8 years) individuals. Specifically, for each region, we randomly sampled 7 males and 7 females (6 for the LGN) and used these individuals to re-create sex prediction models (using both X chromosome and autosomal genes) and estimate known sex prediction probabilities per individual.

#### Sex differences in gene expression heterogeneity

To investigate sex and age differences in gene expression heterogeneity, we calculated Euclidean distances for residual gene expression between all pairs of samples using the euc function in the R package bioDist (108), maintained all pairs of same-sex, same-age group (< or > 8 years), and cross-individual samples, and compared average distances between sex and age group combinations using Tukey’s HSD.

#### Chromosome overrepresentation analysis

We tested for chromosome overrepresentation among sex-biased genes (LFSR < 0.05 in any region) using one-sided Fisher’s Exact tests (fisher.test function in the R package stats; alternative = ‘greater’). We also ran this analysis on male- and female-biased gene sets separately. P-values for each chromosome were adjusted using the Bonferroni correction (p.adjust function in the R package stats) and adjusted p-values less than 0.05 were considered significant.

#### Tissue specificity

For each gene we calculated τ, a measure of tissue specificity using the following formula (64, 94):

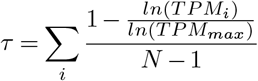

where N is the number of tissues examined, TPMi is the mean TPM per gene within each region, and TPMmax is the highest expression level detected for a given gene over all tissues examined (i.e., maximum TPMi value). The value of τranges from 0 to 1, with lower values indicating an expression pattern that is evenly distributed through all tissues examined and higher τvalues indicating more variation in expression levels across tissues and, thus, a greater degree of tissue specificity. Following other studies (70), we did not normalize the τ calculations in order to reflect this biological reality of gene expression levels. For genes with expression values approaching 0 and low TPMmax, calculations of τ are subject to sampling stochasticity. In order to reduce this effect, TPMi was set to 1 for samples with no detected expression (i.e., less than 1 TPM) (70). Per gene, we modelled log(τ) as a function of the absolute value of the difference between male mean and female mean residual expression values (calculated as in Aim 3) (averaged across all samples). To test for a significant relationship between these variables, we: 1) calculated the Spearman’s rank order correlation; and 2) calculated the moving average log(τ) across non-overlapping window increments of 0.05 expression-level differences, and then calculated the best-fit linear regression line for this moving average. We ran these analyses on all nonY chromosome genes (as their expression is limited to males).

#### Comparisons of loss-of-function (LOF) mutation tolerance

We used LoFtools (110) to assign LOF metrics to each gene. This database is based on the ratio of loss-of-function to synonymous mutations, with lower LoFtool percentiles representing more intolerance to functional variation. Homo sapiens gene names from the LoFtools list were converted to (*Macaca mulatta*) Ensembl gene IDs using the bioMart R package, resulting in a set of 7786 orthologous genes. Per gene, we modeled LOF intolerance percentiles as a function of the absolute value of the difference between male and female mean residual expression values (see above) across all samples and regions. We tested for a significant association between these variables as above (see Tissue specificity).

#### Comparisons of genetic variance

For each gene within each region, the genetic variance of expression (*vu*) was estimated from the EMMA models described above. Per gene and within each region, we modeled log(*vu*) as a function of the absolute value of the difference between male and female mean residual expression values (see above). The structure of this data resulted in a bimodal distribution for estimated values of vu, so we evaluated the relationship between *vu* and sex differences in expression separately within each distribution. We tested for a significant association between these variables as above (see Tissue specificity).

