## Supplementary Materials for "Evolutionary and biomedical implications of sex differences in the primate brain transcriptome"

This file includes:

- Supplementary Text
- Supplementary Figures (including titles and legends)

### Supplementary Text

After adjusting gene expression levels for estimated cell type proportions (Methods), 3.3% (422/12672) of brain expressed genes were differentially expressed in at least one region (Supplementary Table 9), and most of these genes exhibited sex-bias in a majority of regions (58.8% were biased in at least 8 tissues) (Figure 2; Supplementary Figure 7). Of these, 9% were X-linked, 2% were Y-linked, and 89% were autosomal (Supplementary Figure 8). Although the number of male-biased genes was higher than the number of female-biased genes for 14/15 regions (Figure 2), female-biased genes exhibited significantly larger (t-test: $p < 0.001$) magnitudes of their sex effects (mean $|\beta| = 0.10$) than male-biased genes (mean $|\beta| = 0.07$; excluding N = 9 Y chromosome genes, which are not expressed in females). In particular, female-biased X-linked genes exhibited significantly larger sex effects (N=25 genes; mean $|\beta| = 0.20$) than either male-biased X-linked genes (N = 14; mean $|\beta| = 0.10$; Tukey's HSD $p_{adj} < 0.001$) or sex-biased autosomal genes (female: N = 196, mean $|\beta| = 0.08$, $p_{adj} < 0.001$; male: N = 183: mean $|\beta| = 0.07$, $p_{adj} < 0.001$) (Supplementary Figure 8).

Although only 59% of genes identified as sex-biased from cell type-corrected data were also sex-biased in at least one region in our primary analysis, enrichment analyses using either data set tended to produce similar results (Figure 2). Specifically, female-biased genes were associated with translation, cell cycle, apoptosis, and the immune system (T cells, B cells), and male-biased genes were associated with protein metabolism, cell cycle, and vesicular transport (Supplementary Table 16). For the 52 biological processes that were enriched in both analyses, p-values were significantly correlated across data sets for male- and female-biased genes (male: $\rho = 0.740$, $p < 0.001$; female: $\rho = 0.787$, $p < 0.001$) (Supplementary Figure 9). Although we did not detect enrichment for any diseases when using cell type-corrected data (Supplementary Table 17), D values were significantly correlated across data sets for male- and female-biased genes (N=1257 diseases; male: $\rho = 0.722$, $p < 0.001$; female: $\rho = 0.704$, $p < 0.001$) (Supplementary Figure 9). Similarly, we did not detect enrichment of ASD-upregulated or downregulated genes among female- or male-biased gene sets ($p > 0.05$). Cell-type corrected data also produced similar motif enrichments (correlation of ranked p-values: $\rho = 0.548$, $p < 0.001$) (Supplementary Figure 9), including significant enrichment of the estrogen-related receptor alpha motif (Erra: OR = 1.085, p = 0.011) (Supplementary Table 18; Supplementary Figure 9).

### Supplementary Figures

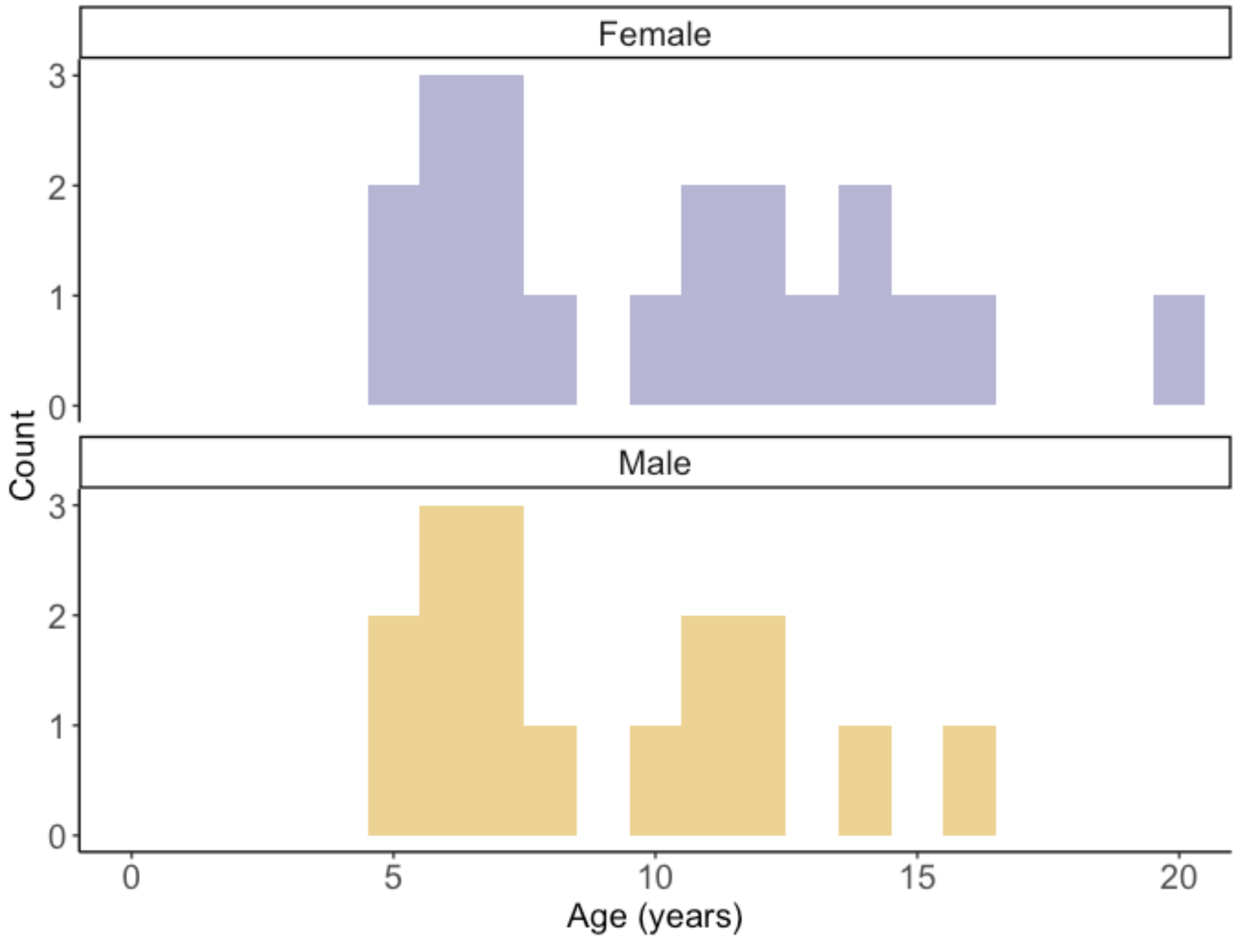


**Figure S1**

Age distribution of adult male and female macaques included in this study.

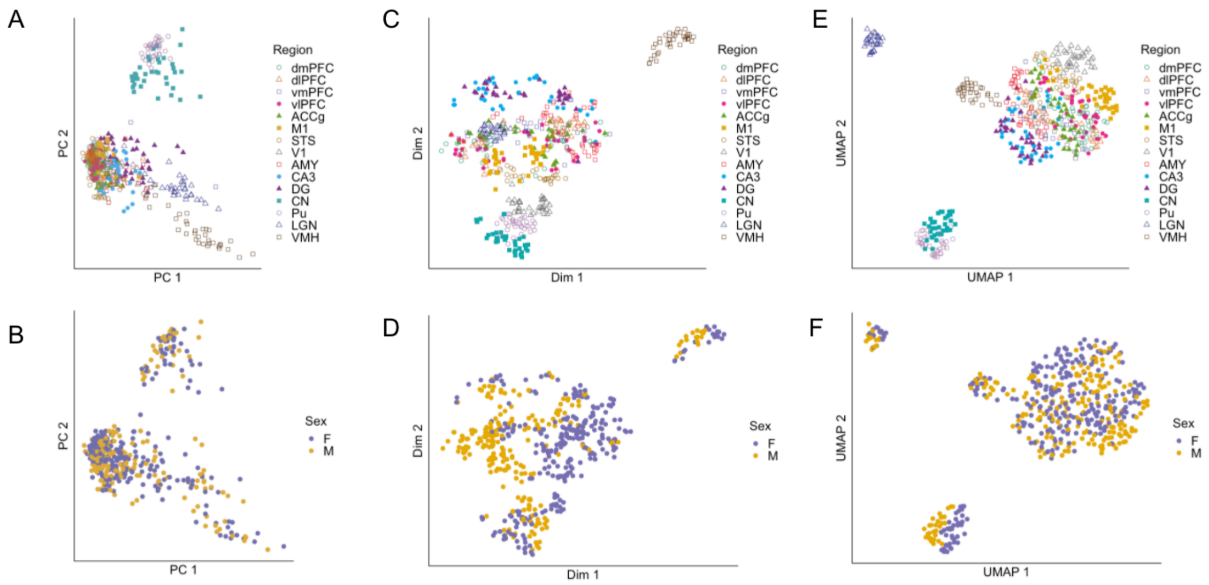


**Figure S2**

A) Principal components analysis (PCA) plot of expression data. Each point represents one sample (N=527). Colors and shapes indicate region (see legend).
B) Principal components analysis (PCA) plot of expression data. Each point represents one sample (N=527). Colors indicate sex (see legend).
C) t-SNE plot of expression data. Each point represents one sample (N=527). Colors and shapes indicate region (see legend).
D) t-SNE plot of expression data. Each point represents one sample (N=527). Colors indicate sex (see legend).
E) Uniform Manifold Approximation and Projection (UMAP) plot of expression data. Each point represents one sample (N=527). Colors and shapes indicate region (see legend).
F) Uniform Manifold Approximation and Projection (UMAP) plot of expression data. Each point represents one sample (N=527). Colors indicate sex (see legend).

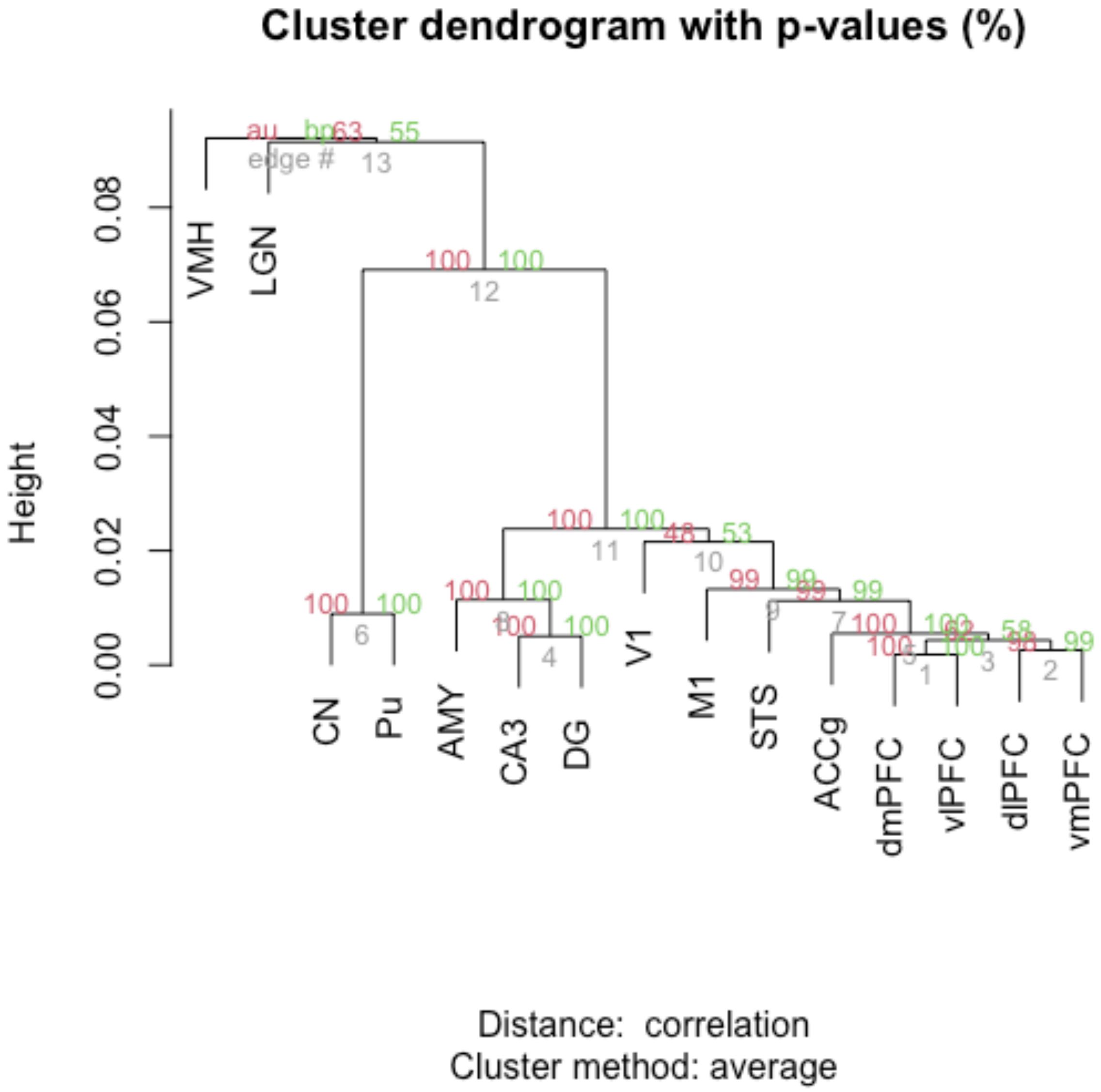


**Figure S3**

Dendrogram depicting hierarchical clustering of brain regions (using expression values averaged across samples within each region after removing batch effects). Approximately unbiased (AU) p-values (red) and bootstrap probability (BP) values (green) are depicted for each cluster. AU values are calculated using multiscale bootstrap resampling, while BP values are calculated by the ordinary bootstrap resampling.

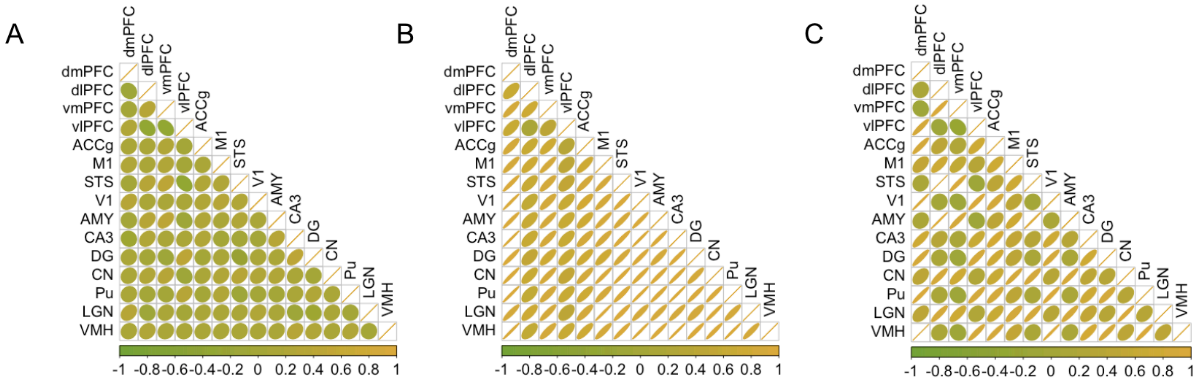


**Figure S4**

Correlation plots. Yellow = positive correlation, green = negative correlation, circle = weak correlation; oval = stronger correlation in the given direction.

A) Correlations are shown for un-shrunken sex effect sizes (from EMMREML) using cell type corrected data across regions. Of these inter-regional correlations, 83 are significantly positive, 15 are significantly negative, and are 7 not significant (p>0.05).

B) Correlations are shown for shrunken sex effect sizes (from MASHR) using cell type corrected data across regions. Of these inter-regional correlations, 105 are significantly positive.

C) Correlations are shown for shrunken sex effect sizes (from MASHR) using cell type corrected data across regions. Of these inter-regional correlations, 96 are significantly positive and 9 are significantly negative.

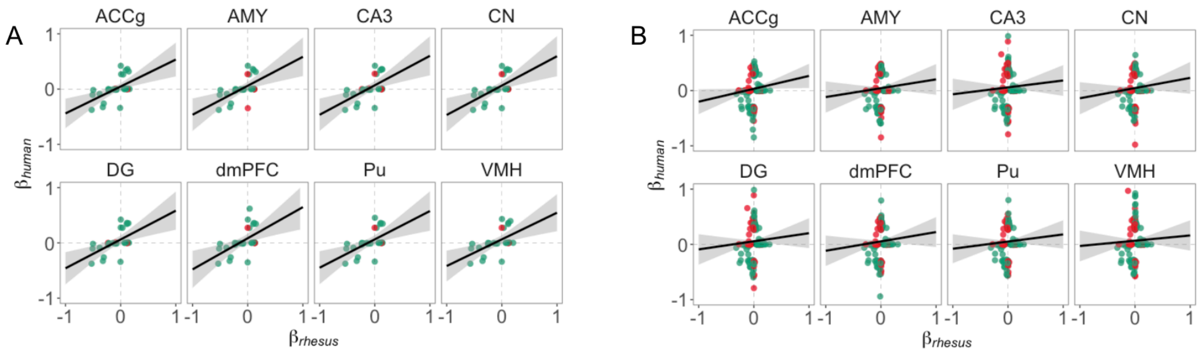


**Figure S5**

A) Scatterplots of estimated sex effects for genes that are significantly sex-biased (LFSR < 0.05) in either humans (GTEx) or rhesus macaques (this study). Only X chromosome genes are included. Green points represent genes with concordant sex-bias across species, while red points represent discordance.
B) Scatterplots of estimated sex effects for genes that are significantly sex-biased (LFSR < 0.05) in either humans (GTEx) or rhesus macaques (this study). Only autosomal genes are included. Green points represent genes with concordant sex-bias across species, while red points represent discordance.

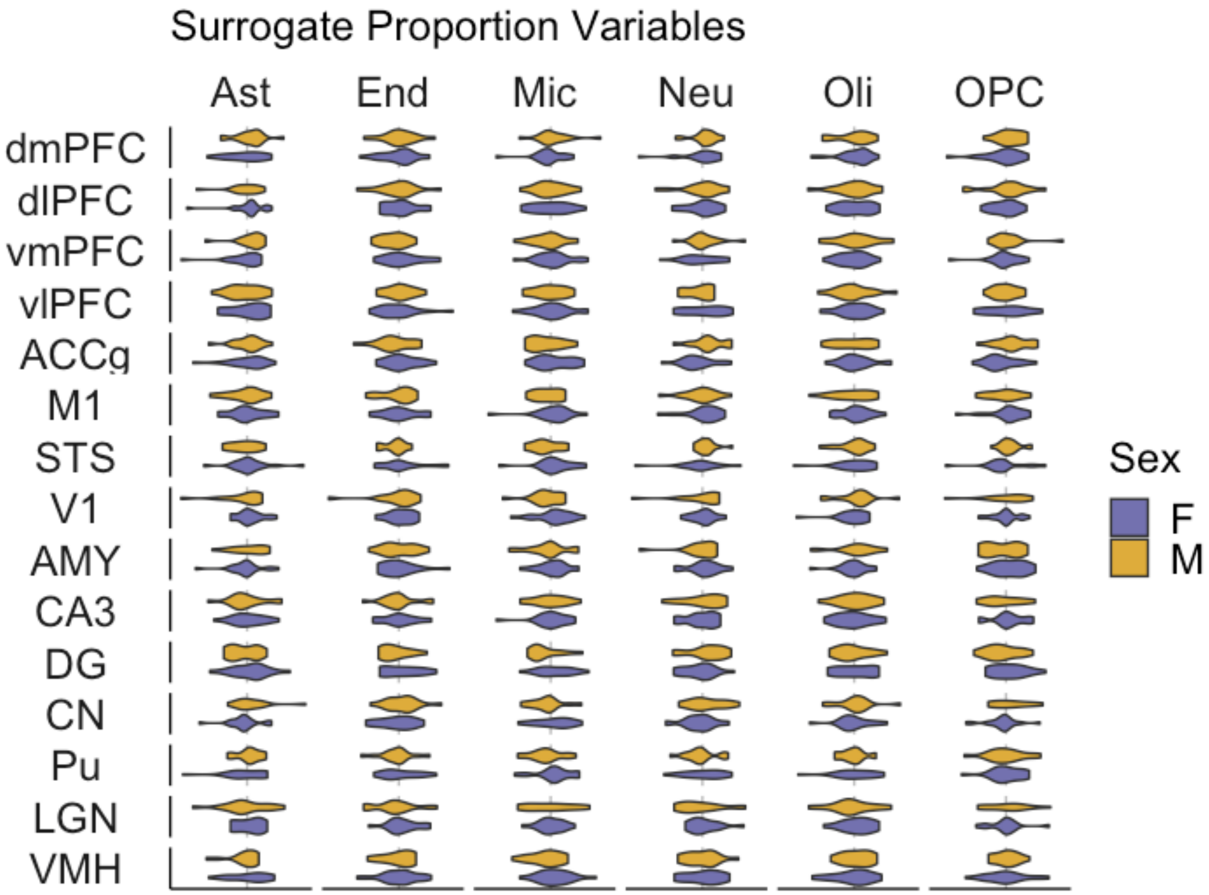


**Figure S6**

Violin plots of estimated surrogate proportion variables for males and females within each region and for each of six brain cell types. Ast = astrocytes; End = endothelial; Mic = microglia; Neu = neurons, Oli = oligodendrocytes; OPC = oligodendrocyte precursor cells

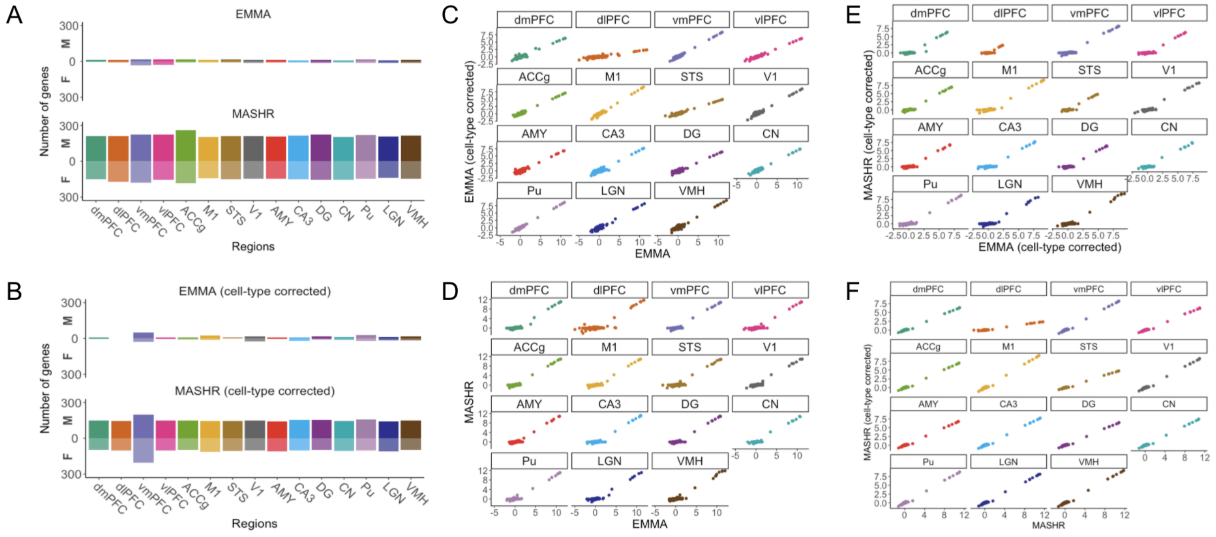


**Figure S7**

A) Counts of sex-biased genes identified using EMMREML (top) and mashr (bottom) on unadjusted expression data. M = male-biased, F = female-biased.
B) Counts of sex-biased genes identified using EMMREML (top) and mashr (bottom) on cell-type adjusted expression data. M = male-biased, F = female-biased.
C) Sex effects (betas) estimated for each gene within each region (x-axis: EMMREML, unadjusted expression data; y-axis: EMMREML, cell-type adjusted expression data)
D) Sex effects (betas) estimated for each gene within each region (x-axis: mashr, unadjusted expression data; y-axis: EMMREML, unadjusted expression data)
E) Sex effects (betas) estimated for each gene within each region (x-axis: EMMREML, cell-type adjusted expression data; y-axis: mashr, cell-type adjusted expression data)
F) Sex effects (betas) estimated for each gene within each region (x-axis: mashr, unadjusted expression data; y-axis: mashr, cell-type adjusted expression data)

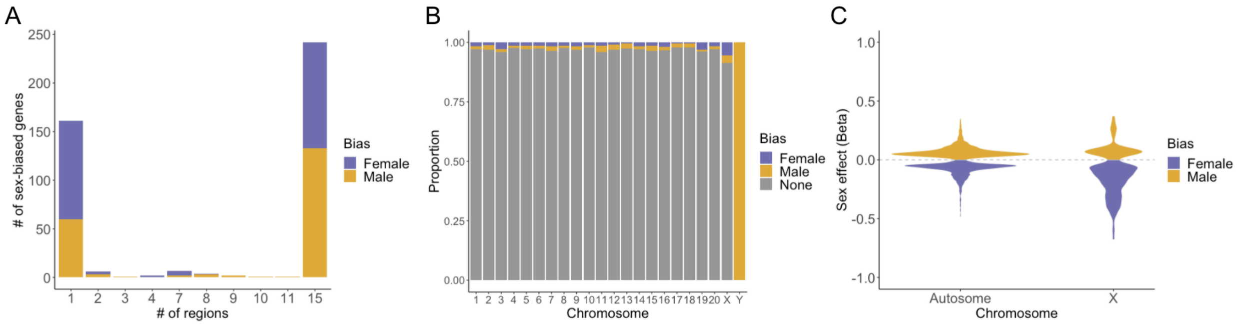


### Figure S8

Results for cell-type adjusted mashr analyses.
A) Stacked bar chart of the number of sex-biased genes shared across different numbers of regions.
B) Proportions of genes on each chromosome that are not biased in any region (grey), female-biased in at least 1 region (purple), or male-biased in at least 1 region (yellow).
C) Violin plots of sex effect sizes for sex-biased autosomal versus X chromosome genes.

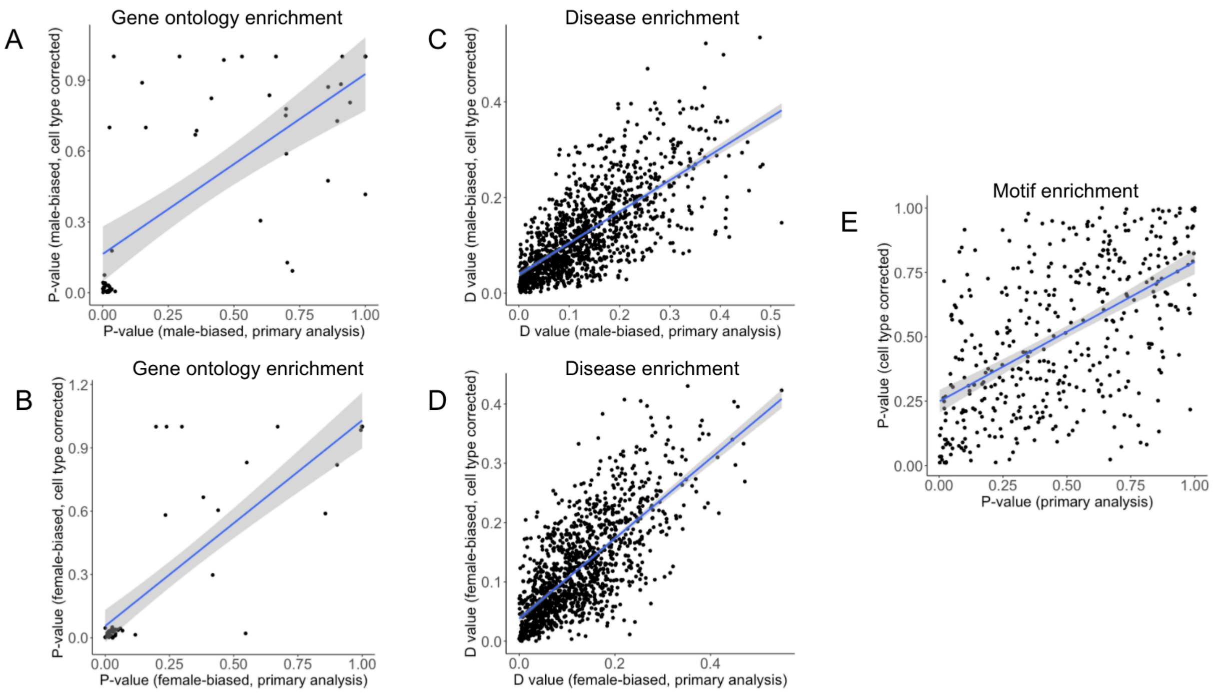


**Figure S9**

A) P-values for N=52 overlapping biological processes across primary and cell-type-corrected analyses (direction = male-biased; $\rho$ = 0.740; p < 3.686e-10)
B) P-values for N=52 overlapping biological processes across primary and cell-type-corrected analyses (direction = female-biased; $\rho$ = 0.787; p < 4.479e-12)
C) D values for N=1257 diseases across primary and cell-type-corrected analyses (direction = male-biased; $\rho$ = 0.722; p < 2.2e-16)
D) D values for N=1257 diseases across primary and cell-type-corrected analyses (direction = female-biased; $\rho$ = 0.704; p < 2.2e-16)
E) P-values for N=414 motifs across primary and cell-type-corrected analyses ($\rho$ = 0.548; p < 2.2e-16)

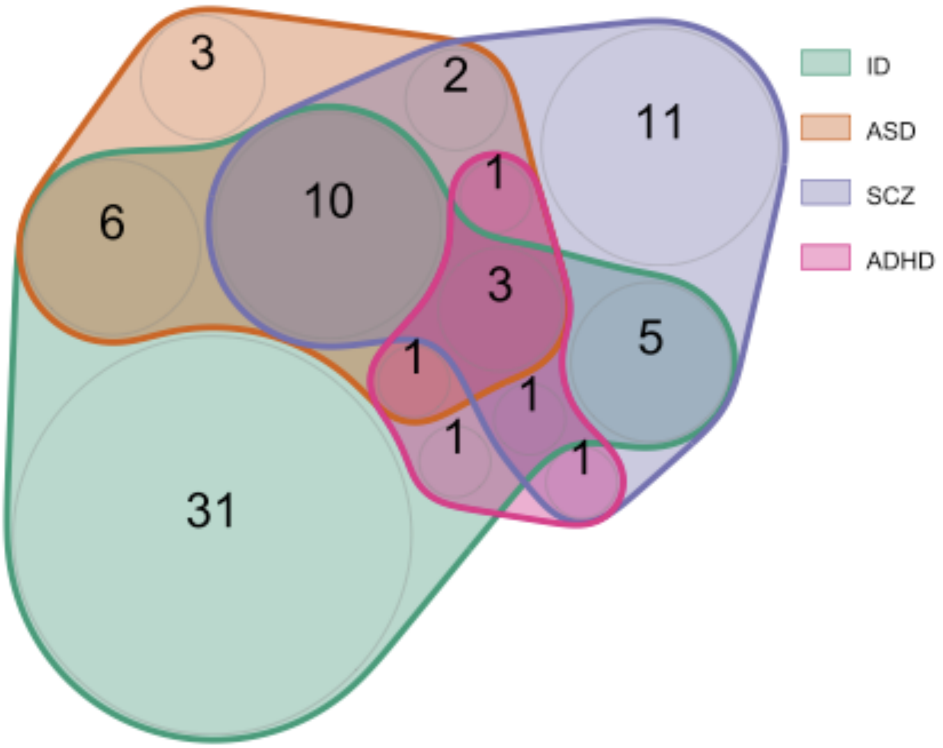


**Figure S10**

Venn diagram of genes associated with each condition that are also male-biased in our dataset (LFSR < 0.2). ID = intellectual disability; ASD = autism spectrum disorder; SCZ = schizophrenia; ADHD = attention deficit hyperactivity disorder.

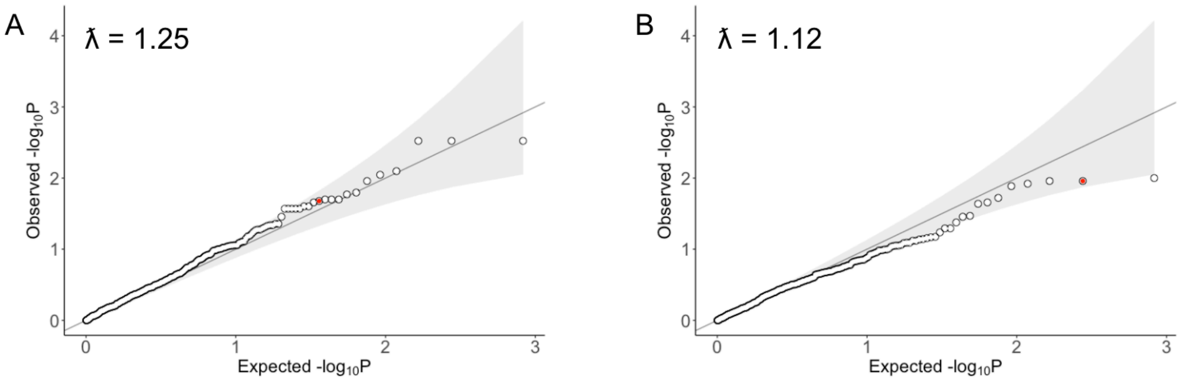


**Figure S11**

A) Observed versus expected p-value distribution for motif enrichment. Red point indicates value for estrogen receptor (Erra). λ̃ = lambda.

A) Observed versus expected p-value distribution for motif enrichment (using cell-type-corrected expression data). Red point indicates value for estrogen receptor (Erra). λ̃ = lambda.

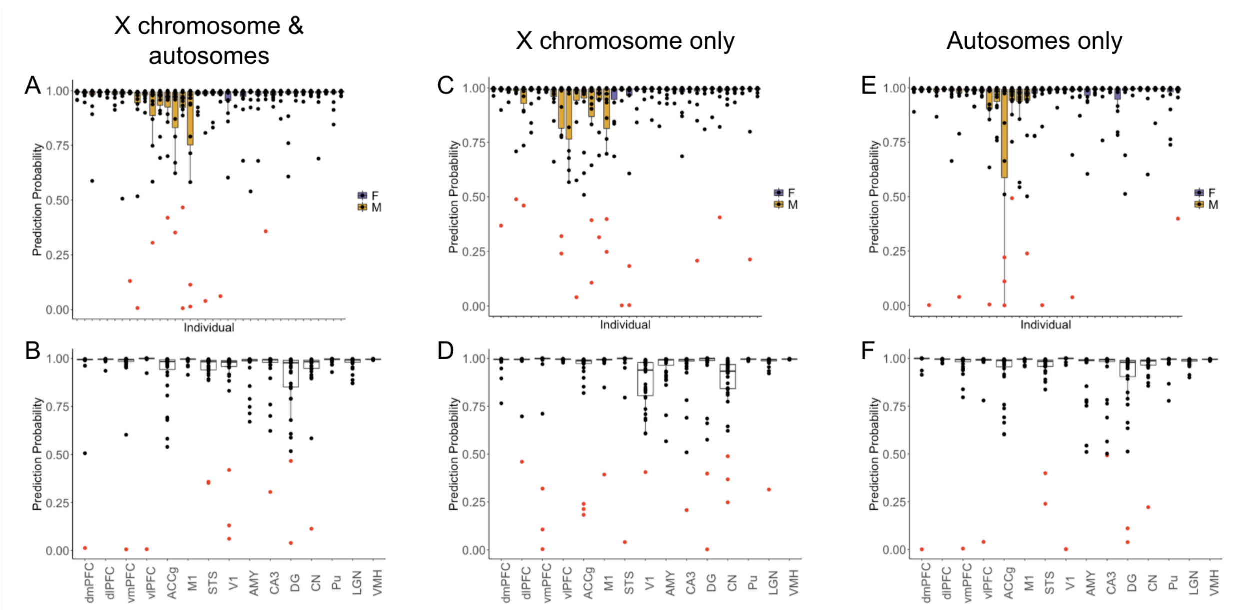


**Figure S12**

A-F) Boxplots of prediction probabilities of the known sex per individual (top row) and region (bottom row). Dots indicate values for individual samples. Purple boxes = female, yellow boxes = male, black dots = correctly classified samples, red dots = incorrectly classified sample (prediction probability of correct sex < 0.5). Results are shown separately for models run incorporating all X chromosome and autosomal genes (A,B), X chromosome genes only (C,D), and autosomal genes only (E,F).

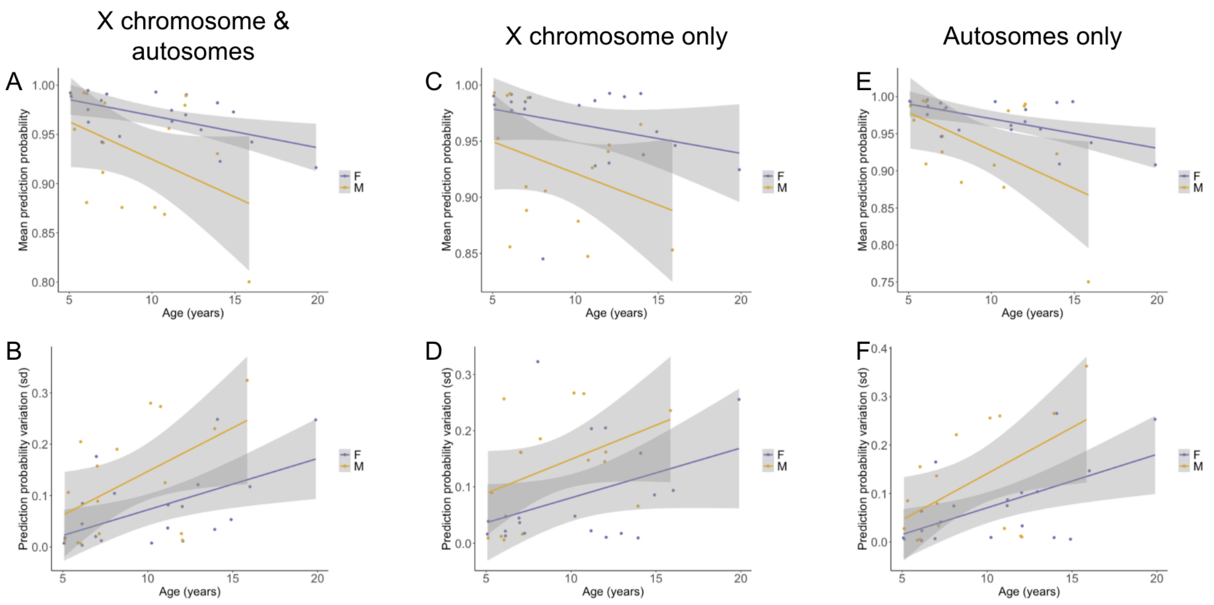


**Figure S13**

A) Average prediction probability of known sex as a function of age (years) for females (purple) and males (yellow) or models run incorporating all X chromosome and autosomal genes (all: estimate = -0.003750, p = 0.0606; females: estimate = -0.003259, p = 0.00798; males: estimate = -0.007654, p = 0.0893)
B) Average standard deviation of prediction probability of known sex as a function of age (years) for females (purple) and males (yellow) or models run incorporating all X chromosome and autosomal genes (all: estimate = 0.010668, p = 0.0089; females: estimate = 0.009981, p = 0.0115; males: estimate = 0.01698, p = 0.0426)
C) Average prediction probability of known sex as a function of age (years) for females (purple) and males (yellow) or models run incorporating X chromosome genes only (all: estimate = -0.002656, p = 0.211; females: estimate = -0.002640, p = 0.202; males: estimate = -0.005653, p = 0.17)
D) Average standard deviation of prediction probability of known sex as a function of age (years) for females (purple) and males (yellow) or models run incorporating X chromosome genes only (all: estimate = 0.008390, p = 0.0469; females: estimate = 0.008853, p = 0.0853; males: estimate = 0.012116, p = 0.0983)
E) Average prediction probability of known sex as a function of age (years) for females (purple) and males (yellow) or models run incorporating autosomal genes only (all: estimate = -0.005151, p = 0.0158; females: estimate = -0.003971, p = 0.00482; males: estimate = -0.010242, p = 0.0356)
F) Average standard deviation of prediction probability of known sex as a function of age (years) for females (purple) and males (yellow) or models run incorporating autosomal genes only (all: estimate = 0.012182, p = 0.00411; females: estimate = 0.011033, p = 0.00782; males: estimate = 0.019074, p = 0.0315)

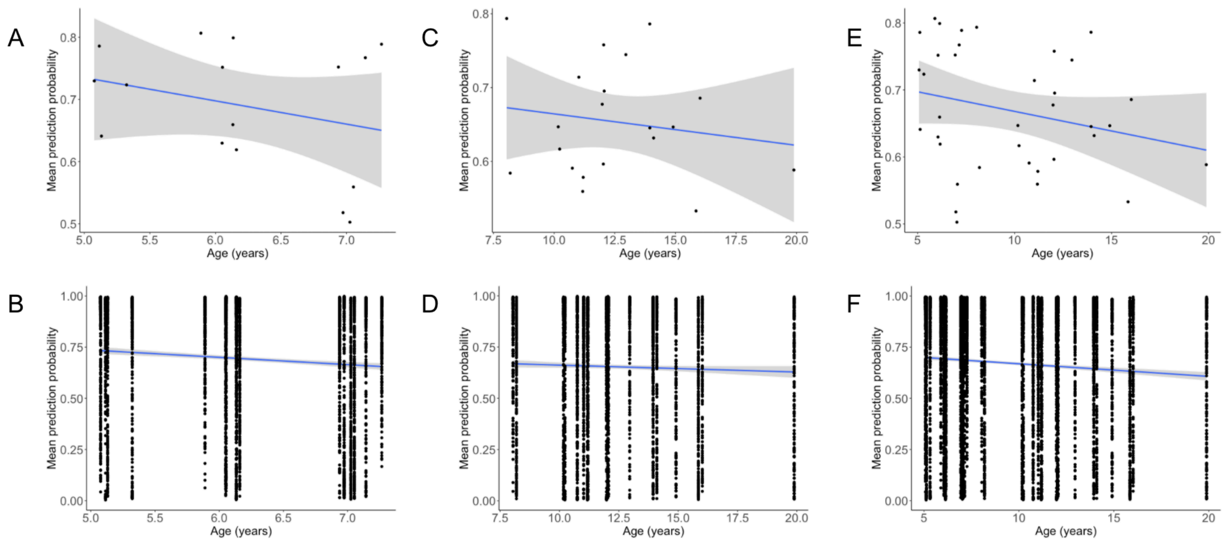


**Figure S14**

A) Prediction probability per individual (<=8 years group), averaged across regions and simulations (estimate = -0.037, p = 0.282).
B) Prediction probability per individual (<=8 years group) for each region and simulation (estimate = -0.003, p = 0.061).
C) Prediction probability per individual (>8 years group), averaged across regions and simulations (estimate = -0.004, p = 0.509).
D) Prediction probability per individual (>8 years group) for each region and simulation (estimate = -0.036, p = 1.08e-08).
E) Prediction probability per individual (combined results from separate analyses of both age groups), averaged across regions and simulations (estimate = -0.006, p = 0.139).
F) Prediction probability per individual (combined results from separate analyses of both age groups) for each region and simulation (estimate = -0.006, p = 2.49e-11).

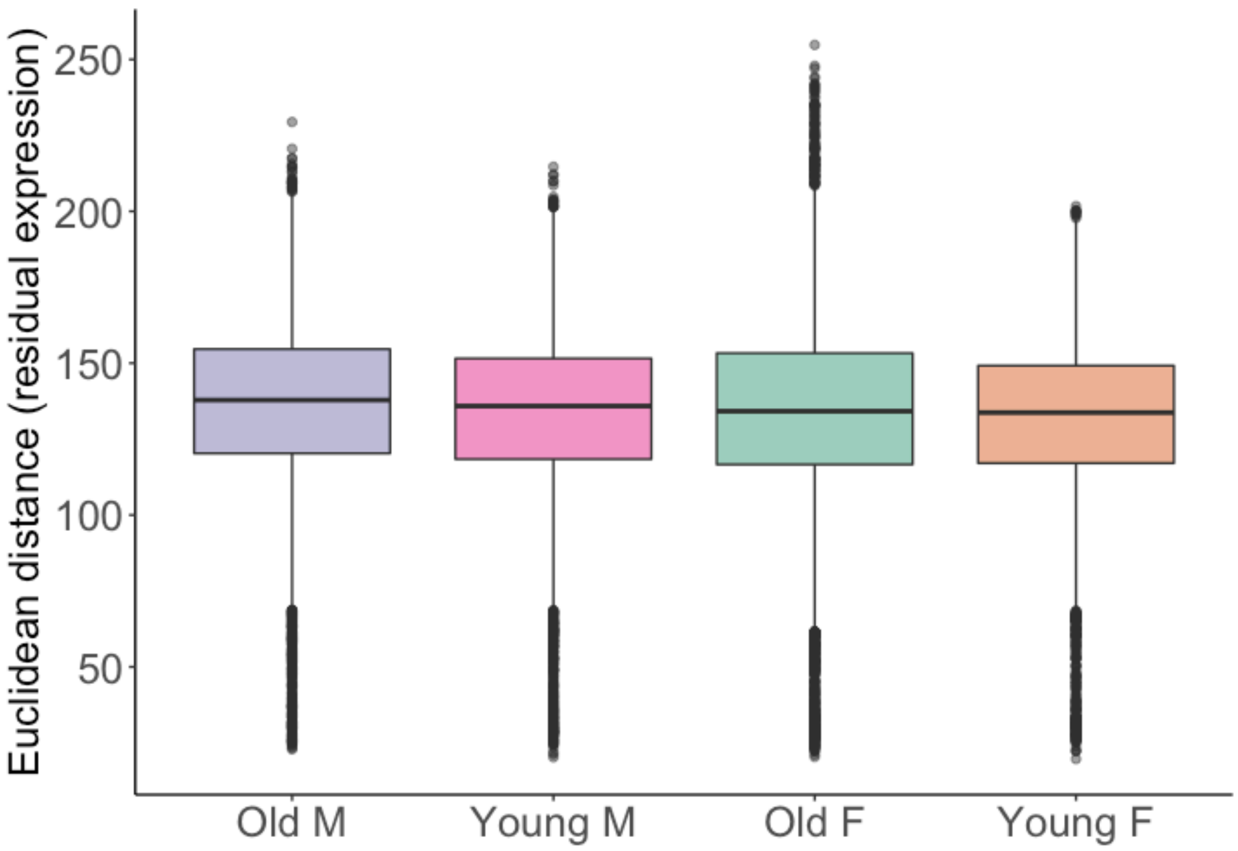


**Figure S15**

M = males; F = females; Old individuals are > 8 years old; Young individuals are < 8 years old. Groups are ordered from highest (left) to lowest (right) median values. Boxplots indicate the median (black horizontal line), first and third quartiles (i.e., interquartile range, IQR; lower and upper hinges), and ranges extending from each to 1.5 x IQR beyond each hinge (whiskers). Points represent individual genes that are outliers (i.e., beyond whiskers).

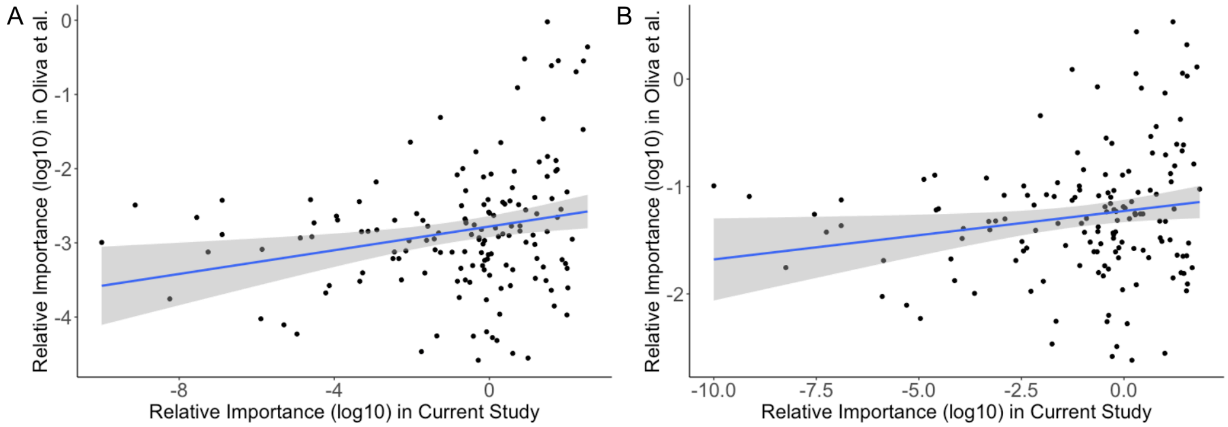


**Figure S16**

Relative importance of X chromosome genes for sex prediction in X chromosome gene models in the current study and Oliva et al. (2020).

A) Relevance is summed across regions in both studies ($\rho$ = 0.222, p = 0.006).

B) Relevance is averaged across regions in both studies ($\rho$ = 0.0170, p = 0.038).

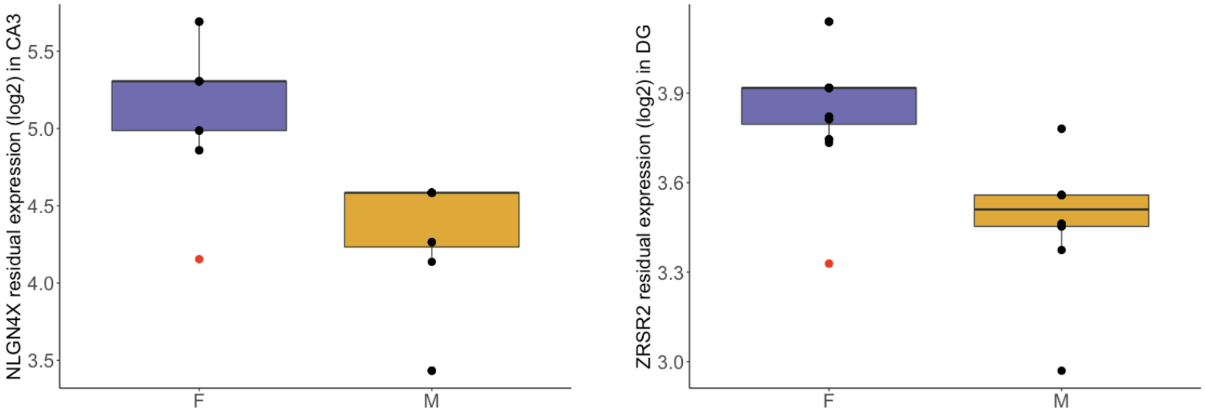


**Figure S17**

Boxplots of the residual expression values for the most influential gene in a given region from X chromosome gene models (left: NLGN4X in CA3; right: ZRSR2 in DG). The red point represents the misclassified female sample. In humans, ZRSR2 escapes and NLGN4X variably escapes XCI. Boxplots indicate the median (black horizontal line), first and third quartiles (i.e., interquartile range, IQR; lower and upper hinges), and ranges extending from each to 1.5 x IQR beyond each hinge (whiskers). Points represent individual genes that are outliers (i.e., beyond whiskers).

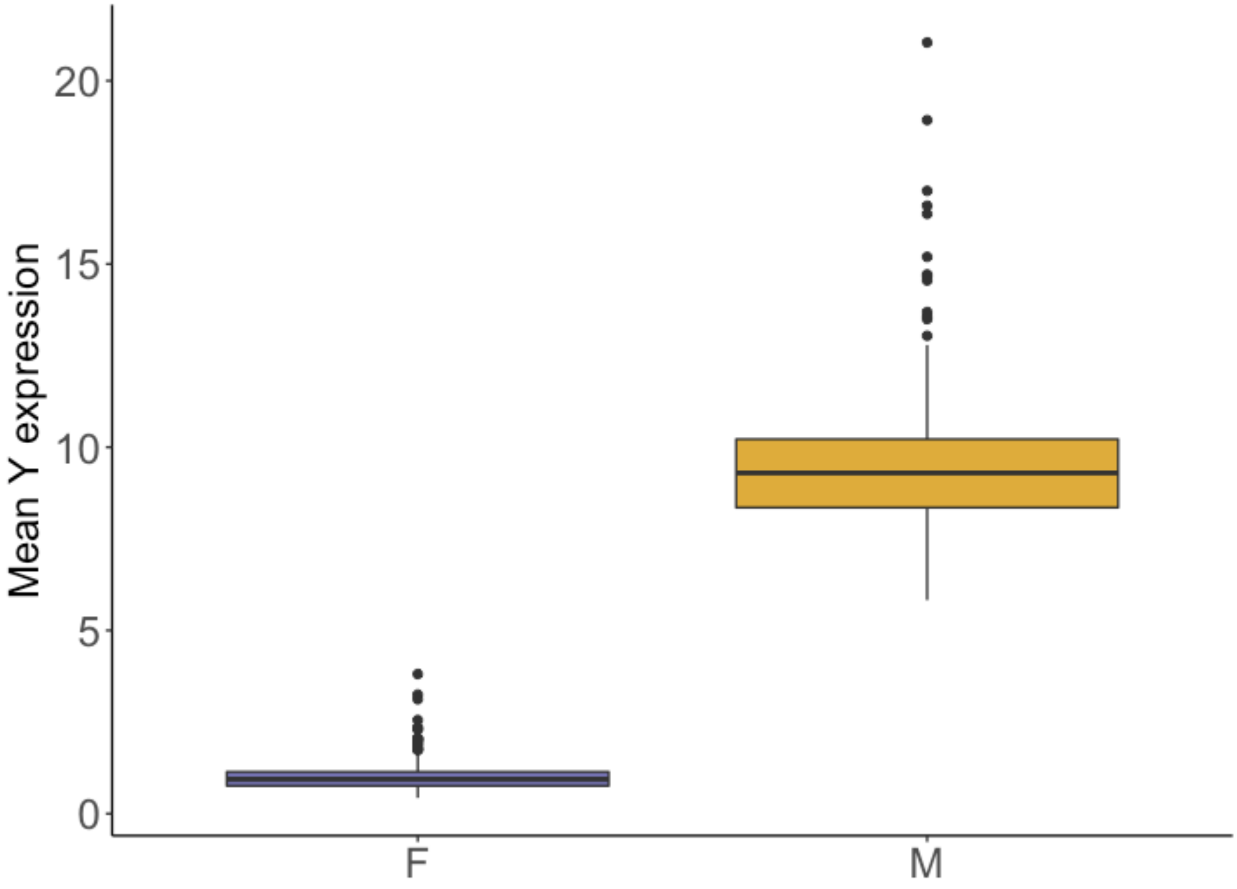


**Figure S18**

Mean expression (TPM) of all Y genes for female and male samples. Reads were mapped to the original (non-sex specific) transcriptome. The female reads detected here represent the gametologues (X-Y homologues) and are not present in the sex-specific mapped data used in the manuscript. Boxplots indicate the median (black horizontal line), first and third quartiles (i.e., interquartile range, IQR; lower and upper hinges), and ranges extending from each to 1.5 x IQR beyond each hinge (whiskers). Points represent individual genes that are outliers (i.e., beyond whiskers).
